# Stress-induced vtRNA1-1 modulates redox homeostasis and ferroptosis susceptibility in hepatocellular carcinoma cells

**DOI:** 10.64898/2026.03.30.715395

**Authors:** EunBin Kong, Daniel Sanchez-Taltavull, Carolina Oliveira Rizzo, Anastasiia Suspitsyna, Deborah Stroka, Norbert Polacek

## Abstract

Ferroptosis is a unique form of regulated cell death characterized by iron–dependent lipid peroxidation. Although the molecular details of ferroptosis regulation have been widely explored, the contributions of short non-coding RNAs (ncRNAs) to ferroptosis regulation, other than miRNAs remain poorly understood. Here, we identified vault RNA1-1 (vtRNA1-1) as a previously unrecognized short ncRNA regulator of ferroptosis in hepatocellular carcinoma (HCC) cells. vtRNA1-1 expression was upregulated by ferroptosis inducers and exhibited strong negative correlation with ferroptosis sensitivity, thus protecting cells from ferroptosis. vtRNA1-1 levels were elevated in selected ferroptosis–resistant cells, while its depletion reversed the phenotype thus resensitizing these cells to ferroptosis. These findings suggested a contribution of vtRNA1-1 to both intrinsic and acquired ferroptosis resistance. Mechanistically, we uncovered that increased oxidative stress, which potentiates lipid peroxidation, specifically induced expression of the vtRNA1-1 paralog in an NF-κB dependent manner. Elevated vtRNA1-1 levels suppressed NF-κB-mediated pro-oxidant gene expression, thereby limiting reactive oxygen species (ROS) accumulation and alleviating oxidative stress. Taken together, oxidative stress–inducible vtRNA1-1 governs redox balance by forming a reciprocal regulatory loop with NF-κB and this loop determines ferroptosis susceptibility by adjusting basal ROS levels. Our findings provide unprecedented insights into the regulation of redox homeostasis in HCC cells mediated by a short ncRNA and uncovered vtRNA1-1 as a potential therapeutic target for overcoming ferroptosis resistance in liver cancer.

## 1. Introduction

Regulating the balance between cell death and survival is a fundamental process for maintaining homeostasis [1]. Multiple regulated cell death (RCD) pathways, including apoptosis, necroptosis, autophagy-dependent cell death, and pyroptosis, coordinate this balance in response to diverse stresses [2]. Among these, ferroptosis has recently emerged as an iron-dependent form of RCD driven by uncontrolled lipid peroxidation [3], with critical implications for cancer progression and therapeutic response [4,5]. Ferroptosis is distinct from other types of cell death from biochemical, genetic, and morphological perspectives. Morphologically, ferroptotic cells exhibit ruptured mitochondria and reduced plasma membrane integrity, without showing distinct changes in the nucleus [6–8]. As a key lipid antioxidant enzyme, glutathione peroxidase 4 (GPX4) plays a pivotal role in protecting cells from ferroptotic cell death by neutralizing oxidized phospholipids [9]. Redox homeostasis, balancing reactive oxygen species (ROS) production and antioxidant defense activation [10], is central for several cell death pathways, including ferroptosis [11,12]. Either accumulation of excessive ROS or a defective cellular antioxidant system induces oxidative stress, thereby triggering ferroptosis through lipid peroxidation [13,14]. In this process, ferrous ions play a central role in generating hydroxyl radicals via the Fenton reaction, leading to uncontrollable lipid peroxidation [15]. Accordingly, maintenance of redox homeostasis is critical to regulate ferroptosis.

Non-coding RNAs (ncRNAs) are a diverse class of transcripts that do not encode proteins. In recent years, ncRNAs have been recognized as essential regulators across a wide range of cellular processes from orchestrating gene expression to fine-tuning signaling pathways in all domains of life [16]. Among the ncRNAs, several types of short ncRNAs such as Y RNA, and tRNA-derived RNA (tDR) have been reported to participate in cellular responses to various stress conditions, thereby contributing to cellular homeostasis [17–21]. Recent studies have uncovered the functions of ncRNAs in maintaining redox homeostasis and regulating ferroptosis [22,23]. Numerous micro RNAs (miRNAs) and long non-coding RNAs (lncRNAs) modulate signaling pathways that govern redox balance, including the NRF2–Keap1 or NF-κB pathways [24,25]. For instance, lncRNA nuclear paraspeckle assembly transcript 1 (NEAT1) competitively binds to miR-362-3p and activates myo-inositol oxygenase (MIOX) expression to promote ferroptosis [26], and circTTC13 inhibits ferroptosis by targeting mir-513a-5p [27]. Additionally, various miRNAs directly target key factors such as GPX4, SLC7A11, NRF2 or GLS2 to regulate redox homeostasis and ferroptosis [28]. However, the functional relevance of short ncRNAs other than miRNAs in redox homeostasis and ferroptosis regulation is largely unknown. Due to their low metabolic cost and rapid turnover rate that provide energetic efficiency and regulatory plasticity, short ncRNAs are considered fast and effective stress-responsive factors in cells. Thus, a deeper understanding of how these short ncRNAs contribute to redox homeostasis may provide novel mechanistic insights.

Vault RNA (vtRNA) is an intriguing class of short ncRNA found in many eukaryotes including humans and mice [29,30]. Human cells express four paralogs of vtRNA (vtRNA1-1, vtRNA1-2, vtRNA1-3, and vtRNA2-1) through RNA polymerase III-dependent transcription [31]. Previous studies have shown that vtRNA1 paralogs play multiple roles in regulating pro-survival characteristics in different immortalized cell lines, which are closely related to tumorigenic features [32–34]. In liver cancer tissues derived from hepatocellular carcinoma (HCC) patients, both vtRNA1-1 and vtRNA1-2 were elevated compared to non-pathological adjacent liver tissues, and their depletion suppresses cancer growth *in cellulo* and in an *in vivo* mouse xenograft model [31,35]. Loss of the vtRNA1-2 paralog impaired critical tumorigenic properties such as cell migration, invasion, proliferation, and angiogenesis [36]. Although its effects on migration and angiogenesis are negligible, vtRNA1-1, on the other hand, promoted the proliferation rates of many human cancer cells and regulates lysosome biogenesis in HCC cells [35]. Furthermore, vtRNA1-1, but not vtRNA1-2, inhibits apoptosis under several stress conditions, including starvation and viral infection, in various human cancer cell lines [31–33]. These findings indicate that each vtRNA1 paralog enhances cell survivability and tumorigenicity in distinct manners. However, the relationship between vtRNA and other types of RCD remains entirely unexplored. Given that vtRNA1-1, despite inhibiting apoptosis in most other cancer cell lines, showed no association with apoptosis in the HCC cell line Huh7 [35], we set out investigating the potential function of vtRNAs in regulating ferroptosis. This goal appeared particularly relevant as the liver is a metabolically active organ generating substantial ROS, potentially leading to various disorders [37].

Here, we uncovered previously unrecognized roles for vtRNA1-1 in maintaining redox homeostasis and conferring resistance to ferroptosis in HCC cells. As an NF-κB–dependent, oxidative stress–inducible ncRNA, vtRNA1-1 negatively correlated with ferroptosis sensitivity and protected liver cells from ferroptosis. We also revealed that this short ncRNA suppresses NF-κB activity and inhibits NF-κB–dependent pro-oxidant expression, thereby controlling basal intracellular ROS levels to maintain redox homeostasis. Given that ferroptosis has emerged as a promising target in various cancers including HCC [38,39], our investigation of the novel vtRNA1-1-mediated regulatory function in governing redox balance and determining ferroptosis susceptibility makes this ncRNA a valuable candidate for therapeutic interventions.

## 2. Materials and methods

### 2.1. Cell culture, treatment, and transfection

The human liver cell lines (Huh7 WT and vtRNA1-1KO, vtRNA1-1 OE, IHH, HepG2, Hep3B, SNU423, SNU445, and SNU475) were cultured in DMEM/F-12 (Gibco, 21331046) supplemented with 10% FBS (Gibco, 10082147) and Pen-Strep Glutamine (Gibco, 10378016). Erastin (MedChemExpress, HY-15763), RSL3 (MedChemExpress, HY-100218A), N-acetylcysteine (MedChemExpress, HY-B0215), Liproxstatin-1 (MedChemExpress, HY-12726), TPCA-1 (MedChemExpress, HY-10074), hydrogen peroxide (Carl Roth, 8070.2), tert-butyl hydroperoxide (Sigma-Aldrich, 458139), dihexyl phthalate (MedChemExpress, HY-W011215), sodium arsenite (Supelco, 1.06277.1000), and Sorafenib tosylate (MedChemExpress, HY-10201A) were used at the indicated concentrations. For reverse transfection of LNA, Lipofectamine RNAiMAX (Invitrogen, 13778075) was used according to the manufacturer’s recommendations. For plasmid transfection, Lipofectamine 3000 (Invitrogen, L3000015) was used according to the manufacturer’s recommendations. For genome editing, CRISPR/Cas9 was used as described in Bracher et al. [34].

### 2.2. Establishment of ferroptosis resistance (FR) cells

Establishment of Ferroptosis resistant (FR) cells was performed as described in Xu et al. [40], by exposing parental Huh7, IHH, and Hep3B cells to stepwise increasing concentrations of Erastin. The resistant phenotype of FR cells was confirmed using CCK-8 assays after treating ferroptosis inducers. FR cells were cultured long-term in DMEM medium supplemented with 2 μM Erastin to maintain the resistant phenotype.

### 2.3. Immunoblot

Cells were harvested and lysed with lysis buffer [20 mM Tris-Cl pH 7.5, 150 mM NaCl, 2.5 mM MgCl_2_, 1% Triton X-100 (Sigma-Aldrich, T8787), 0.25% Na-deoxycholate (Sigma-Aldrich, D6750)], containing protease inhibitor cocktail (Sigma-Aldrich, 11697498001) and phosphatase inhibitor (Thermo Scientific, A32957) for 30 minutes on ice. The supernatants were collected after centrifugation at 16,000 ×g for 10 minutes at 4 °C. Protein concentrations were determined by BCA assay (Thermo Scientific, 23225), and lysates were subsequently boiled in 4× NuPage sample buffer (Invitrogen, NP0007). The protein samples were separated by SDS-PAGE gel and transferred to a 0.45 µm nitrocellulose membrane for immunoblotting. Immunoblotting was performed following standard protocols using the indicated antibodies. All quantified immunoblot signals were normalized to the band intensities of the GAPDH.

### 2.4. Nuclear fractionation

Nuclear fractionation was performed as described in Kong et al. [41]. Cells were harvested and lysed with hypotonic lysis buffer [10 mM Tris-Cl pH 7.5, 10 mM NaCl, 2 mM MgCl_2_], containing protease inhibitor cocktail (Sigma-Aldrich, 11697498001) and phosphatase inhibitor (Thermo Scientific, A32957) for 10 min on ice. The cytosolic fraction supernatants were collected by centrifugation at 1,000 ×g for 10 minutes at 4 °C. Nuclear fraction pellets were washed by PBS with 0.1% Triton X-100 (Sigma-Aldrich, T8787) and collected by centrifugation at 1,000 ×g for 10 minutes at 4 °C. Washed pellets were resuspended by lysis buffer [20 mM Tris-Cl pH 7.5, 150 mM NaCl, 2.5 mM MgCl_2_, 1% Triton X-100 (Sigma-Aldrich, T8787), 0.25% Na-deoxycholate (Sigma-Aldrich, D6750)], containing protease inhibitor cocktail (Sigma-Aldrich, 11697498001) and phosphatase inhibitor (Thermo Scientific, A32957) and retained by sonication, twice on ice.

### 2.5. RNA extraction, reverse transcription-PCR (RT-PCR), and quantitative real-time PCR (qPCR)

Total RNA was extracted from the harvested cells using TRI-reagent (Zymo Research, R2050-1-200) according to the manufacturer’s recommendations. Reverse transcription of total RNA was performed using SuperScript IV One-Step RT-PCR System (Invitrogen, 18090010), and random hexamers following the manufacturer’s recommendations. qPCR reactions were performed using Rotor Gene-Q (QIAGEN), and analyses were conducted in Robotics software (QIAGEN) according to the manufacturer’s instructions.

### 2.6. Northern blot

For northern blot, 4-6 µg of extracted total RNA was separated on a denaturing polyacrylamide gel (7 M Urea, 1x TBE buffer), transferred to a nylon membrane (Amersham Hybond N+; GE Healthcare, RPN203B), UV-crosslinked, and probed with 5′-^32^P-end labeled antisense DNA probes as described in Gebetsberger et al. [42]. Signals were exposed to a phosphor screen and quantified using a Typhoon FLA1000 phosphorimager and ImageQuantTL software. All quantified northern blot signals were normalized to the band intensities of the 5.8S rRNA or EtBr-stained 5S rRNA.

### 2.7. Cell viability assay

Cell viability was measured by CCK-8 (MedChemExpress, HY-K0301). After incubating cells with indicated nucleic acids or drug at different time point, cell viability was assessed following the manufacturer’s recommendations. The absorbance was measured at 450 nm using a Tecan Infinite M1000Pro plate reader.

### 2.8. Detection of intracellular reactive oxygen species (ROS)

Intracellular ROS was measured by DCFH-DA (MedChemExpress, HY-D0940). After treatment with 50 µM H_2_O_2_ for 30 minutes and washing with PBS, cells were incubated with 20 µM DCFH-DA for 30 minutes in the dark and washed by PBS three times, followed by adding a live cell imaging solution (Invitrogen, A59688DJ). Images were captured using a fluorescence microscope using LAS X software (Leica Microsystems, DMI6000B) according to the manufacturer’s recommendations.

### 2.9. Detection of lipid peroxidation

Lipid peroxidation was measured by BODIPY 581/591 C11 (MedChemExpress, HY-D1301). Cells were treated with the indicated ferroptosis inducers or inhibitor for 2.5 hours and washed with PBS. In the presence of the same drugs, cells were incubated with 10 µM of BODIPY 581/591 for 30 minutes in the dark and washed by PBS three times, followed by adding a live cell imaging solution (Invitrogen, A59688DJ). Images were captured using a fluorescence microscope using LAS X software (Leica Microsystems, DMI6000B) according to the manufacturer’s recommendations.

### 2.10. Bioinformatic analysis

Transcriptomic analysis was performed on mRNAs sequencing data from wild-type and vtRNA1-1 knockout Huh7 cells (Gene Expression Omnibus, GSE276534) using DESeq2 [36]. Dysregulated genes with adjusted p-value < 0.05 and log2 fold change ≥ 1 in vtRNA1-1 knockout vs WT were defined as the vtRNA1-1-associated signature. TCGA-LIHC primary tumor transcriptomic data was analyzed using the TCGA biolinks R package. Expression values were extracted as TPM and transformed as log2(TPM + 1). Signature scores present in TCGA-LIHC dataset were standardized across samples using a z-score. For each sample, the signature score was defined as the mean z-score across signature genes. Samples were stratified into Top20 and Bottom80 groups using the 80^th^ percentile of the score distribution. A list of ferroptosis-related genes was obtained from the FerrDB [43], and lists of NRF2 target genes, NF-kB target genes, and oxidative stress responsive genes were obtained from the GSEA and the Molecular Signatures Database (MSigDB) [44,45]. Gene symbols were mapped to Ensembl identifiers using TCGA annotation. For each geneset, genes present in the TCGA-LIHC were selected and standardized by gene-wise z-score across samples and scores of each sample were calculated as the mean z-score across genes. For visualization, scores were scaled to a 0–100 range within each geneset. Heatmaps were generated from expression matrices using the R package pheatmap. Boxplots and PCA plots were generated using ggplot2 and custom R scripts. PCA was performed on log2(TPM + 1) expression matrix using prcomp.

## 3. Results

### 3.1 vtRNA1-1 levels negatively correlate with ferroptosis sensitivity in liver cells

vtRNA1-1 has been considered a pro-survival factor that inhibits apoptosis in many cancer cell lines, such as HeLa, BL41, BL2, A549, HEK293 and Hs578T [32,33]. However, in the HCC cell line Huh7, altered vtRNA1-1 levels had no significant effect on apoptosis [35]. Thus, we examined the relationship between vtRNA1-1 and one of the most prominent non-apoptotic cell death pathways in HCC cells, ferroptosis. To investigate their relationship, we re-analyzed our published transcriptome data [36] which revealed significant changes of the transcriptional landscapes of ferroptosis-related genes between vtRNA1-1 knockout (KO) cells and wild-type (WT) control cells **(Fig. 1A)**. To further define this potential association, we treated Huh7 cells with the ferroptosis inducers RSL3 and Erastin and assessed vtRNA levels. Ferroptosis induction upregulated vtRNA1-1 but did not affect vtRNA1-2 or vtRNA1-3 expression **(Fig. 1B, 1C, S1A and S1B)**, supporting a potential relationship between vtRNA1-1 and ferroptosis.

**Figure 1.**
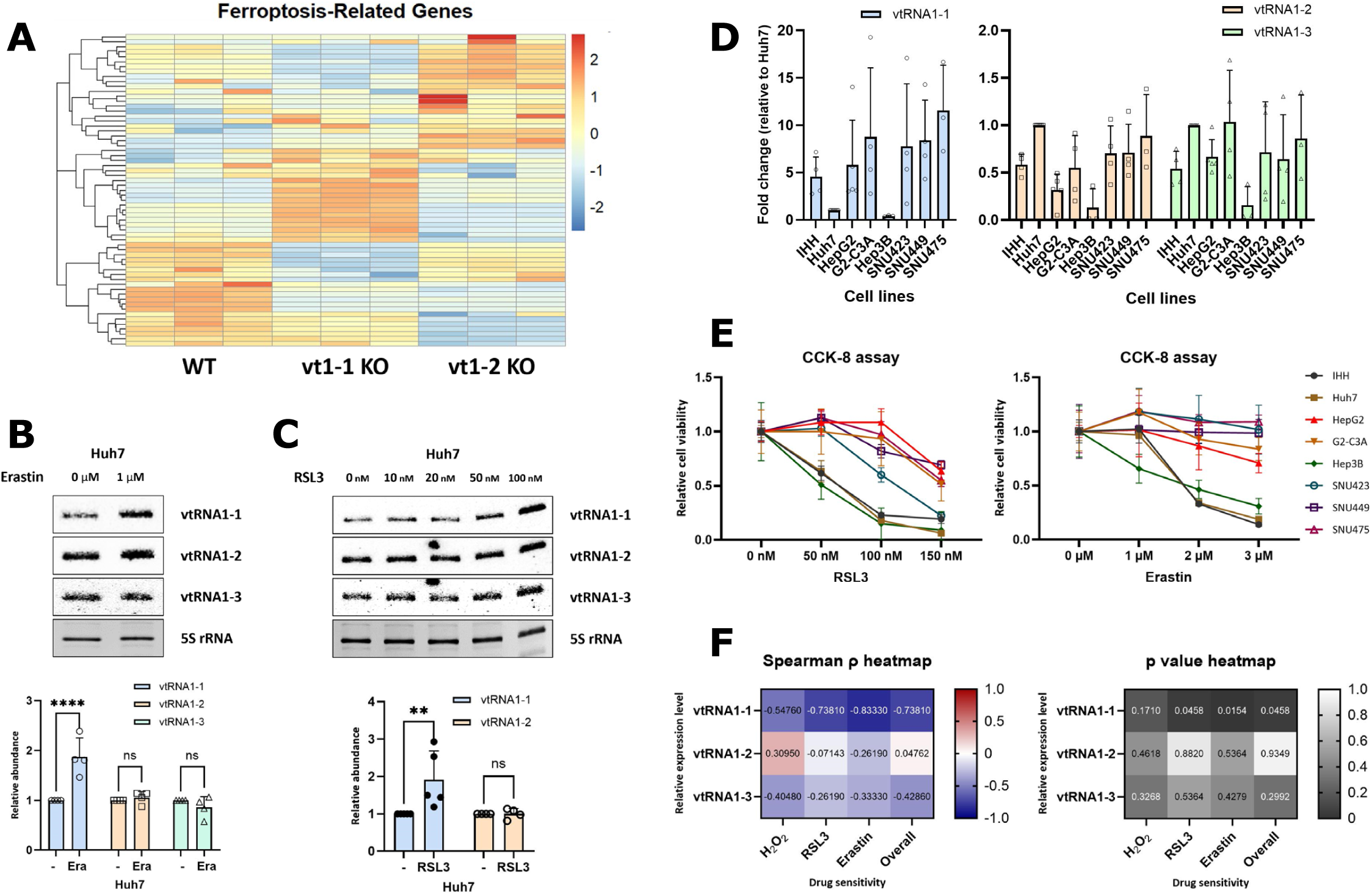
vtRNA1-1 expression shows a strong negative correlation with ferroptosis sensitivity. (A) A heatmap showing the expression levels of ferroptosis-related genes in Huh7 WT, vtRNA1-1 KO, and vtRNA1-2 KO cells. (B and C) Northern blot analyses of vtRNAs. Total RNA was extracted from Huh7 cells treated with indicated ferroptosis inducers for 24 hours and analyzed. 5S rRNA served as a loading control. The intensity of vtRNAs was normalized to 5S rRNA (bottom, n ≥4). Error bars indicate the standard deviation. ****p<0.0001, **p<0.01, ns: non-significant (two-way ANOVA test) (D) Quantification of vtRNA1 paralogs across eight human liver cell lines. Northern blot intensities were normalized to 5S rRNA (n≥3), and relative expression of each vtRNA1 paralog in different cell lines is presented as fold change normalized to Huh7 cells. Representative blot is shown in figure S1C. (E) The indicated cells were treated with RLS3 (left) and Erastin (right) for 24 hours and the cell viability was measured by CCK-8 assay. (F) Heatmaps showing Spearman’s correlation coefficient (ρ, left) and p-value (right) between each vtRNA1 paralog expression and sensitivity to different ferroptosis inducers. The color indicates Spearman’s correlation coefficient (left) or p-value (right). |ρ|≥0.7: very strong, 0.7>|p|≥0.4: strong, 0.4>|p|≥0.3: moderate, 0.3>|p|≥0.2: weak, 0.2>|p|≥0: negligible relationship

We then conducted a comparative analysis using eight different liver cell lines, including IHH, Huh7, HepG2, HepG2-C3A, Hep3B, SNU423, SNU449, and SNU475, to gain a deeper insight into their relationship. The levels of vtRNA1 paralogs were assessed in each liver cell line **(Fig. 1D and S1C)**. Interestingly, despite originating from the same *VTRNA-1* locus on chromosome 5, all three vtRNA1 paralogs displayed markedly different expression patterns across the cell lines investigated. The vtRNA1-1 paralog was most abundant in IHH, HepG2, HepG2-C3A, SNU423, SNU449 and SNU475 cells, whereas in Huh7 cells vtRNA1-2 showed the strongest expression. Notably, Hep3B cells possess the lowest expression levels of all three vtRNA1 paralogs. Subsequently, sensitivities to ferroptosis-inducing agents, including RSL3, Erastin and H_2_O_2_, were measured in the same set of cell lines **(Fig. 1E and S1D)**. HepG2, HepG2-C3A, SNU423, SNU449 and SNU475 with relatively high vtRNA1-1 level were resistant to ferroptosis, while Huh7, IHH and Hep3B cell lines were sensitive. Based on these results, we analyzed the correlations between sensitivity to each ferroptosis inducer and expression level of each vtRNA1 paralog (**Fig. 1F**). Spearman’s rank correlation analysis demonstrated that vtRNA1-1 expression showed a very strong negative and statistically significant correlation (Spearman’s ρ≤-0.7) with sensitivity to ferroptosis inducers (RSL3, Erastin, H_2_O_2_ and overall, **Fig. 1F)** On the other hand, vtRNA1-2 and vtRNA1-3 exhibited only weak or negligible correlations with sensitivity to ferroptosis-inducing agents (**Fig. 1F**) In summary, ferroptosis inducers specifically upregulated the vtRNA1-1 paralog, and these expression levels negatively correlated with ferroptosis sensitivity in various liver cell lines.

### 3.2. vtRNA1-1 protects liver cells from ferroptosis

We next investigated the functional role of vtRNA1-1 in ferroptosis. The eight liver cell lines used for correlation analysis were divided into two groups: ferroptosis-sensitive group (Huh7, IHH and Hep3B) and ferroptosis-resistant group (HepG2, G2-C3A, SNU423, SNU449 and SNU475). Cell viability of the former group in response to ferroptosis inducer was assessed after overexpressing vtRNA1-1 or vtRNA1-2. The results revealed that overexpression of vtRNA1-1, but not vtRNA1-2, conferred ferroptosis resistance **(Fig. 2A-C and S2A)**. Conversely, LNA-mediated vtRNA1-1 knockdown in HepG2 and SNU423, which are included in the ferroptosis-resistant group, increased susceptibility to RSL3 and Erastin **(Fig. 2D, 2E and S2B)**. Additionally, reintroduction of vtRNA1-1 into the vtRNA1-1 KO cells recovered cell viability under ferroptotic conditions **(Fig. 2F, S2C and S2D).** Co-treatment with the lipid peroxidation inhibitor Lip-1 rescued cell death, whereas apoptosis inhibitor Z-VAD showed no significant effect **(Fig. S2E)**, corroborating our findings. These data demonstrated that vtRNA1-1 functions as an anti-ferroptosis factor in hepatocytes.

**Figure 2.**
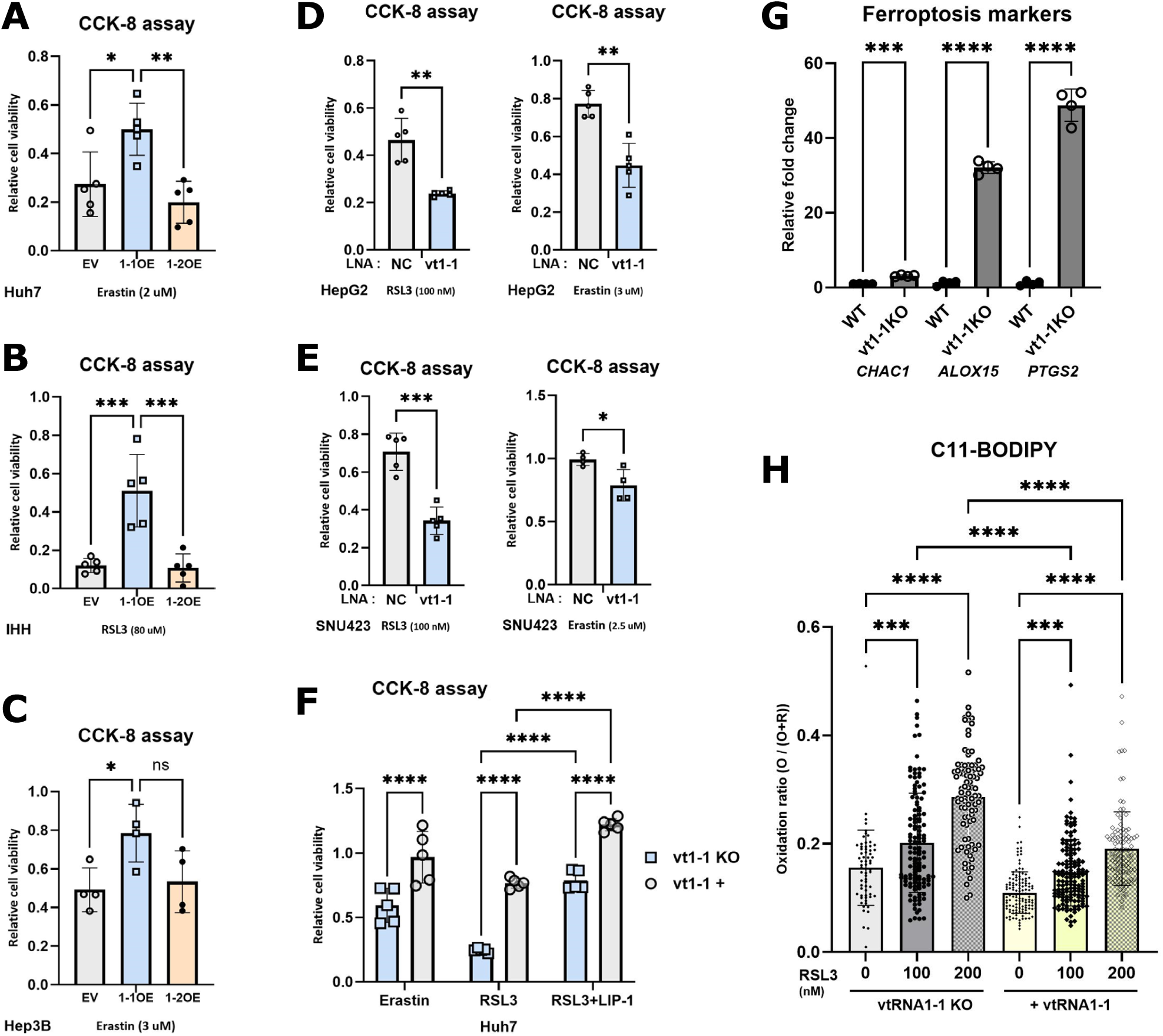
vtRNA1-1 suppresses ferroptotic cell death in hepatocytes. (A-C) Following vtRNA1-1 or vtRNA1-2 overexpression, the indicated cells were treated with ferroptosis inducer for 24 hours, and the relative cell viability was measured by CCK-8 assay, normalized to the untreated cells. EV denotes empty vector (mock transfection). (D and E) Following LNA-mediated knockdown of vtRNA1-1, the indicated cells were treated with ferroptosis inducer for 24 hours, and the relative cell viability was measured by CCK-8 assay, normalized to the untreated cells. NC denotes negative control (mock transfection). (F) vtRNA1-1 KO and vtRNA1-1-expressing (vt1-1 +) Huh7 cells were treated with the indicated drugs for 24 hours, and the cell viability was measured by CCK-8 assay, normalized to the untreated cells. (G) qPCR analysis of the indicated mRNAs in Huh7 WT and vtRNA1-1 KO cells (n=4). (H) Assessment of lipid peroxidation using C11-BODIPY fluorescent probe. vtRNA1-1 KO and vtRNA1-1-expressing (+vtRNA1-1) Huh7 cells were pre-treated with RSL3, and lipid peroxidation was measured. ****p<0.0001, *** p<0.001, **p<0.01, *p<0.05, ns: non-significant (two-way ANOVA test)

Moreover, we observed dysregulated expression of ferroptosis markers in vtRNA1-1 KO cells. CHAC1, ALOX15 and PTGS2 were upregulated in the absence of vtRNA1-1 **(Fig. 2G**), and this dysregulation was significantly increased upon treatment with ferroptosis inducers **(Fig. S2F and S2G),** suggesting that vtRNA1-1 KO cells were more vulnerable to ferroptosis. The intracellular lipid peroxidation assay using a C11-BODIPY fluorescent probe further supported our results that vtRNA1-1 was critical for ferroptosis susceptibility in liver cells. Upon RSL3 treatment, oxidized lipid ratio was elevated in vtRNA1-1 KO cells compared to cells re-expressing vtRNA1-1 **(Fig. 2H and S2H)**, suggesting that vtRNA1-1 exerts a protective effect against ferroptosis.

### 3.3. Depleting vtRNA1-1 overcomes ferroptosis resistance in HCC cells

To determine the impact of vtRNA1-1 in the context of acquired resistance to ferroptosis, we selected ferroptosis-resistant (FR) cells. Parental cells, which were originally placed into the ferroptosis-sensitive group, were exposed to stepwise increasing concentrations of Erastin over several weeks **(Fig. 3A)** [40]. Resistant phenotypes of the established FR cells were verified by measuring cell viability under ferroptotic conditions in all cell lines **(Fig. 3B and S3A)**. The FR cells showed resistance not only to Erastin, but also to other inducers, demonstrating that these cells acquired broad resistance to ferroptosis **(Fig. 3B and S3B)**. Of note, during the selection phase the FR cells did not acquire resistance to sorafenib, a widely used first line drug for HCC **(Fig. S3C)**. To further validate this phenotype, we performed the lipid peroxidation assay, which revealed a smaller increase in oxidized lipid levels upon RSL3 treatment in FR cells compared with parental cells **(Fig. 3C and S3D)**.

**Figure 3.**
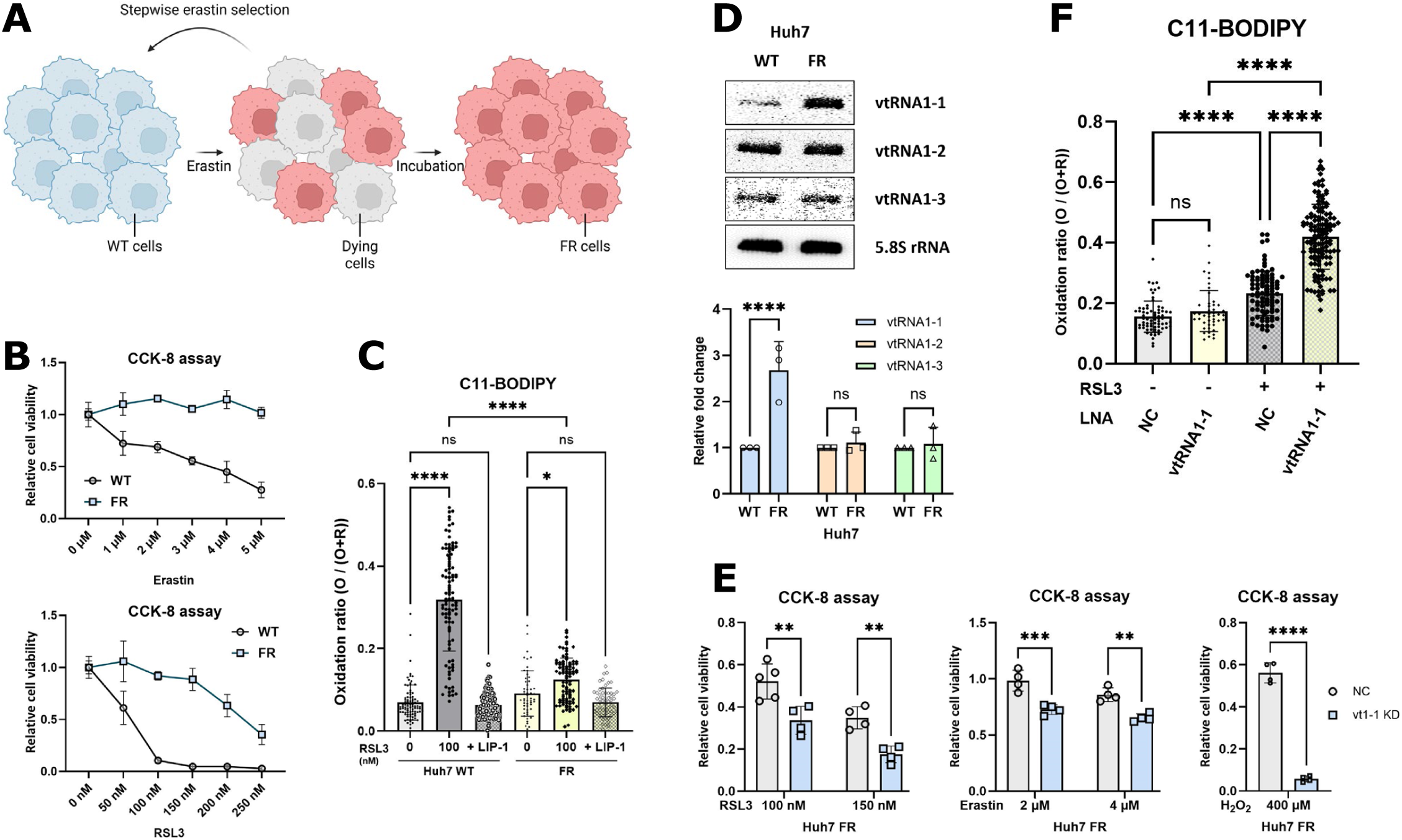
Depletion of vtRNA1-1 re-sensitizes ferroptosis-resistance cells. (A) Establishment of ferroptosis-resistant (FR) cells. Parental cells were selected through stepwise Erastin treatment. (B) Huh7 WT and FR cells were treated with Erastin (Top) and RSL3 (bottom) for 24 hours, and the cell viability was measured by CCK-8 assay. (C) Assessment of lipid peroxidation using C11-BODIPY fluorescent probe. Huh7 WT and FR cells were pre-treated with the indicated drugs, and lipid peroxidation was measured. (D) Northern blot analysis of vtRNAs. Total RNA was extracted from Huh7 WT and FR cells and analyzed. 5.8S rRNA served as a loading control. The intensity of vtRNAs was normalized to 5.8S rRNA (bottom, n=4). Error bars indicate the standard deviation. ****p<0.0001, ns: non-significant (two-way ANOVA test) (E) Following LNA-mediated knockdown of vtRNA1-1, Huh7 FR cells were treated with ferroptosis inducer for 24 hours, and the cell viability was measured by CCK-8 assay, normalized to the untreated cells. NC denotes negative control (mock transfection). (F) Assessment of lipid peroxidation using C11-BODIPY fluorescent probe. Following LNA-mediated knockdown of vtRNA1-1, Huh7 FR cells were pre-treated with RSL3, and lipid peroxidation was measured. ****p<0.0001, *** p<0.001, **p<0.01, *p<0.05, ns: non-significant (two-way ANOVA test)

We subsequently assessed vtRNA1 paralog levels in the established FR cells and found that vtRNA1-1 was markedly upregulated, whereas vtRNA1-2 and vtRNA1-3 showed no changes **(Fig. 3D and S4A).** These results suggested a potential direct involvement of vtRNA1-1 in the acquired ferroptosis resistance. To test this putative causality, we subsequently knocked down the increased vtRNA1-1 levels in the FR cells using locked nucleic acid (LNA) gapmers, resulting in the loss of acquired ferroptosis resistance in the FR cells **(Fig. 3E, S4B-S4D)**. CRISPR-mediated reduction of vtRNA1-1 level in FR cells corroborated this re-sensitization phenotype **(Fig. S4E and S4F)**. Furthermore, the oxidized lipid ratio of the FR cells upon RSL3 treatment was markedly increased by vtRNA1-1 knockdown, revealing a causal role of vtRNA1-1 in the acquired resistant phenotype **(Fig. 3F and S4G)**. As a result, loss of vtRNA1-1 re-sensitized FR cells to ferroptosis, suggesting that vtRNA1-1 could be a potentially effective target to overcome ferroptosis resistance in HCC cells.

### 3.4. Oxidative stress induces vtRNA1-1 in an NF-kB–dependent manner

Given the functional role of vtRNA1-1 in ferroptosis resistance identified above, understanding how this ncRNA is induced under ferroptotic conditions is of interest. To investigate the mechanism by which vtRNA1-1 was upregulated, we first co-treated a lipid peroxidation inhibitor liproxstatin-1 (Lip-1) and an antioxidant N-acetylcysteine (NAC) with ferroptosis inducers. Interestingly, NAC reversed the elevated vtRNA1-1 expression under ferroptotic conditions **(Fig. 4A and S5A)**, while Lip-1 exhibited negligible effects **(Fig. 4A and S5B)**. These findings indicated that vtRNA1-1 upregulation is unlikely to be a direct consequence of lipid peroxidation or subsequent ferroptotic cell death, but is rather dependent on accumulated reactive oxygen species (ROS) in cells, which is commonly observed upon ferroptosis inducer treatment [46–48]. To test this hypothesis, cells were treated with hydrogen peroxide (H_2_O_2_) and vtRNA levels were assessed. Treatment with H_2_O_2_ selectively triggered vtRNA1-1 expression but had no effects on vtRNA1-2 or vtRNA1-3, and this effect was reversed by NAC co-treatment **(Fig. 4B and S5C)**. In line with previous observations, inhibition of lipid peroxidation by Lip-1 treatment could not reverse the effect of H_2_O_2_ on vtRNA1-1 upregulation **(Fig. S5D)**. In addition, treating intracellular ROS-inducing agents, such as tert-butyl hydroperoxide (t-BHP), dihexyl phthalate (DHP), and low concentrations of sodium arsenite, produced similar effects on vtRNA1-1 induction **(Fig. 4C, S5E and S5F)**. We subsequently corroborated that oxidative stress-induced vtRNA1-1 expression was not cell-line specific **(Fig. 4D, S4G, S5H and S5I)**. Overall, our data suggested that oxidative stress mediated by excessive ROS induced vtRNA1-1 in liver cells.

**Figure 4.**
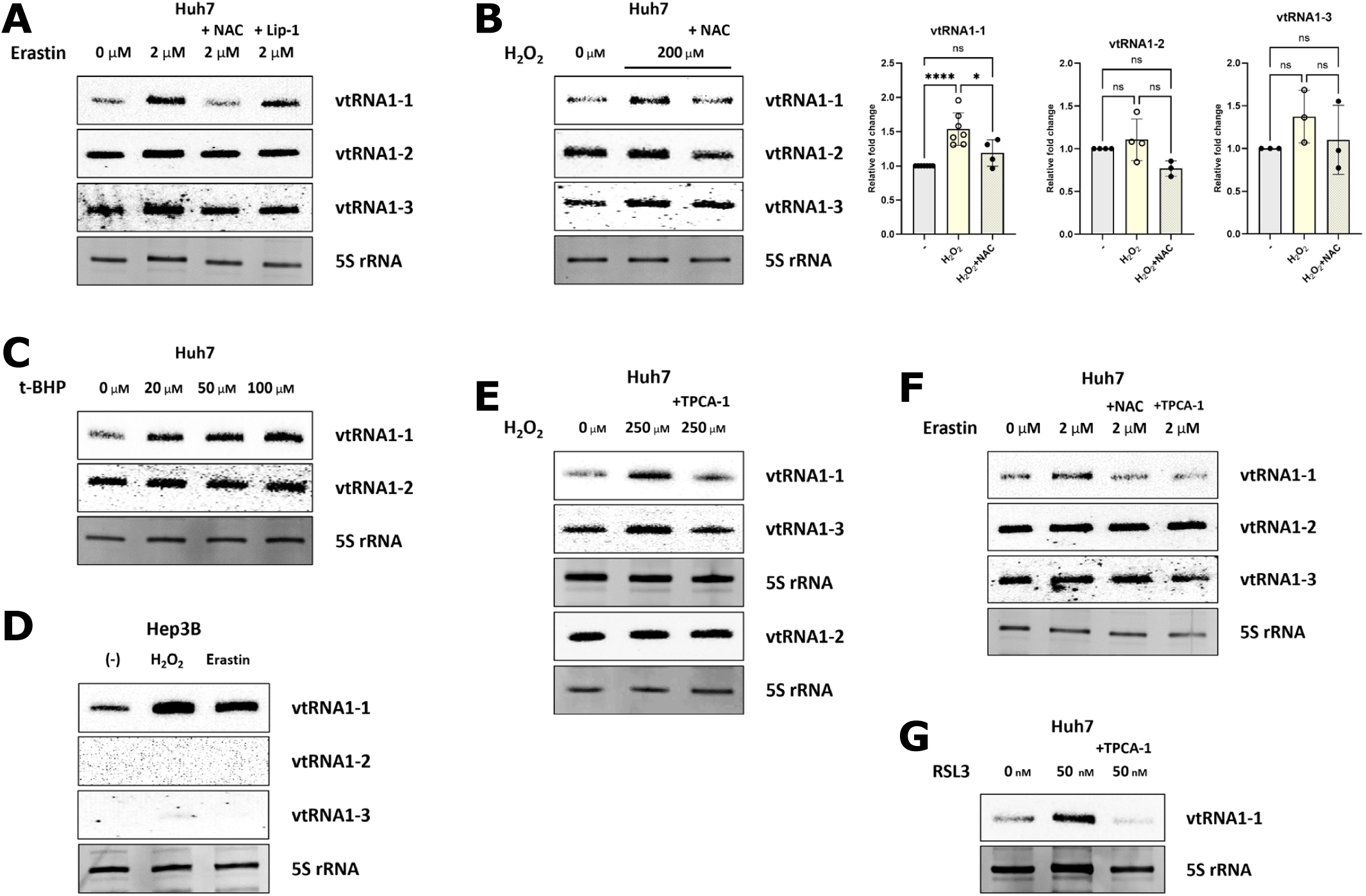
Oxidative stress induces vtRNA1-1 expression via the NF-kB pathway. (A and C-G) Northern blot analyses of vtRNAs. Total RNA was extracted from cells treated with indicated drugs for 24 hours and analyzed. 5S rRNA served as a loading control. Final concentrations were as follows: 5 mM NAC, 1 μM LIP-1, and 10 μM TPCA-1. (B) Northern blot analysis of vtRNAs. Total RNA was extracted from cells treated with indicated drugs for 24 hours and analyzed. 5S rRNA served as a loading control. The intensity of vtRNAs was normalized to 5S rRNA (bottom, n ≥4). Error bars indicate the standard deviation. ****p<0.0001, *p<0.05, ns: non-significant (one-way ANOVA test)

We previously demonstrated that vtRNA1-1 expression is dependent on the NF-κB pathway in several cell lines, including HeLa, HS578T, and BL2. The NF-κB chromatin immunoprecipitation (ChIP) experiments in that study revealed enrichment of NF-κB at the vtRNA1-1 promoter but not at the vtRNA1-2 promoter upon viral infection, indicating that NF-κB is a responsible transcription factor for vtRNA1-1 [32]. Here we verified these findings in liver cells, showing that an NF-κB inhibitor decreased vtRNA1-1 expression, whereas vtRNA1-2 and vtRNA1-3 levels were not affected **(Fig. S4J and S5K)**. Despite the importance of NF-κB signaling on the classical inflammation process, NF-κB is also regarded as a key regulator of the oxidative stress response and is activated by excessive ROS [49,50]. Thus, we examined the role of the NF-κB pathway in vtRNA1-1 upregulation under oxidative stress. Co-treatment of the NF-κB inhibitor TPCA-1 with H_2_O_2_ reversed the upregulation of vtRNA1-1 **(Fig. 4E)**, suggesting that NF-κB upregulates vtRNA1-1 under oxidative stress. This result was further supported by the observation that NF-κB pathway inhibition attenuated the upregulation of vtRNA1-1 by ferroptosis inducers. **(Fig. 4F and 4G)**. Taken together, excessive ROS-mediated oxidative stress activated NF-κB signaling, leading to increased vtRNA1-1 expression, which contributed to ferroptosis resistance in hepatocytes.

### 3.5. vtRNA1-1 attenuates intracellular ROS accumulation by suppressing NF-kB-mediated pro-oxidant expression

To identify how elevated vtRNA1-1 mediates ferroptosis resistance, we first evaluated the expression level of GPX4, a key ferroptosis suppressor [9], but observed no significant changes in vtRNA1-1 KO cells **(Fig. S6A).** Thus, we focused on oxidative stress, a critical driver of ferroptosis. Transcriptional landscape analysis revealed that the expression patterns of oxidative stress responsive genes were clearly distinguishable between vtRNA1-1 KO cells and WT control cells **(Fig. 5A)**. We also examined the expression profile of NF-κB target genes as they play central roles in regulating oxidative stress response and observed significant dysregulation in vtRNA1-1 KO cells **(Fig. 5B)**. This analysis implied that NF-κB is not only a transcription factor responsible for the vtRNA1-1 expression, but also a downstream effector whose activity is regulated by vtRNA1-1. To test this reciprocal regulation between NF-κB and vtRNA1-1, the expression levels of NF-κB-mediated pro-oxidant genes, including DUOX2, NOX1, NOS2, XDH, and PTGS2, were assessed by RT-qPCR. Compared with WT cells, the expression of the pro-oxidants significantly increased in vtRNA1-1 KO cells **(Fig. 5C**), and reintroduction of vtRNA1-1 into KO cells rescued the phenotype **(Fig. S6B)**. Conversely, vtRNA1-1 overexpression suppressed the expression of these genes **(Fig. 5D)**. To corroborate these observations, we examined the nuclear localization of the NF-κB subunit p65, a key indicator of NF-κB activation. In line with our findings, nuclear p65 levels were increased upon vtRNA1-1 depletion **(Fig. 5E)**. Additionally, the phosphorylation of p65 at serine 549 was enhanced in vtRNA1-1 KO cells, suggesting that vtRNA1-1 deficiency activated the NF-κB pathway **(Fig. 5F)**. Moreover, the elevated expression of NF-κB-mediated pro-oxidant genes in vtRNA1-1 KO cells was attenuated by NF-κB inhibition with TPCA-1 **(Fig. 5G)**, highlighting that vtRNA1-1 regulates the pro-oxidants level via NF-κB pathway. Based on these findings, NF-κB and vtRNA1-1 appear to form a reciprocal regulatory loop to modulate the expression of the pro-oxidants.

**Figure 5.**
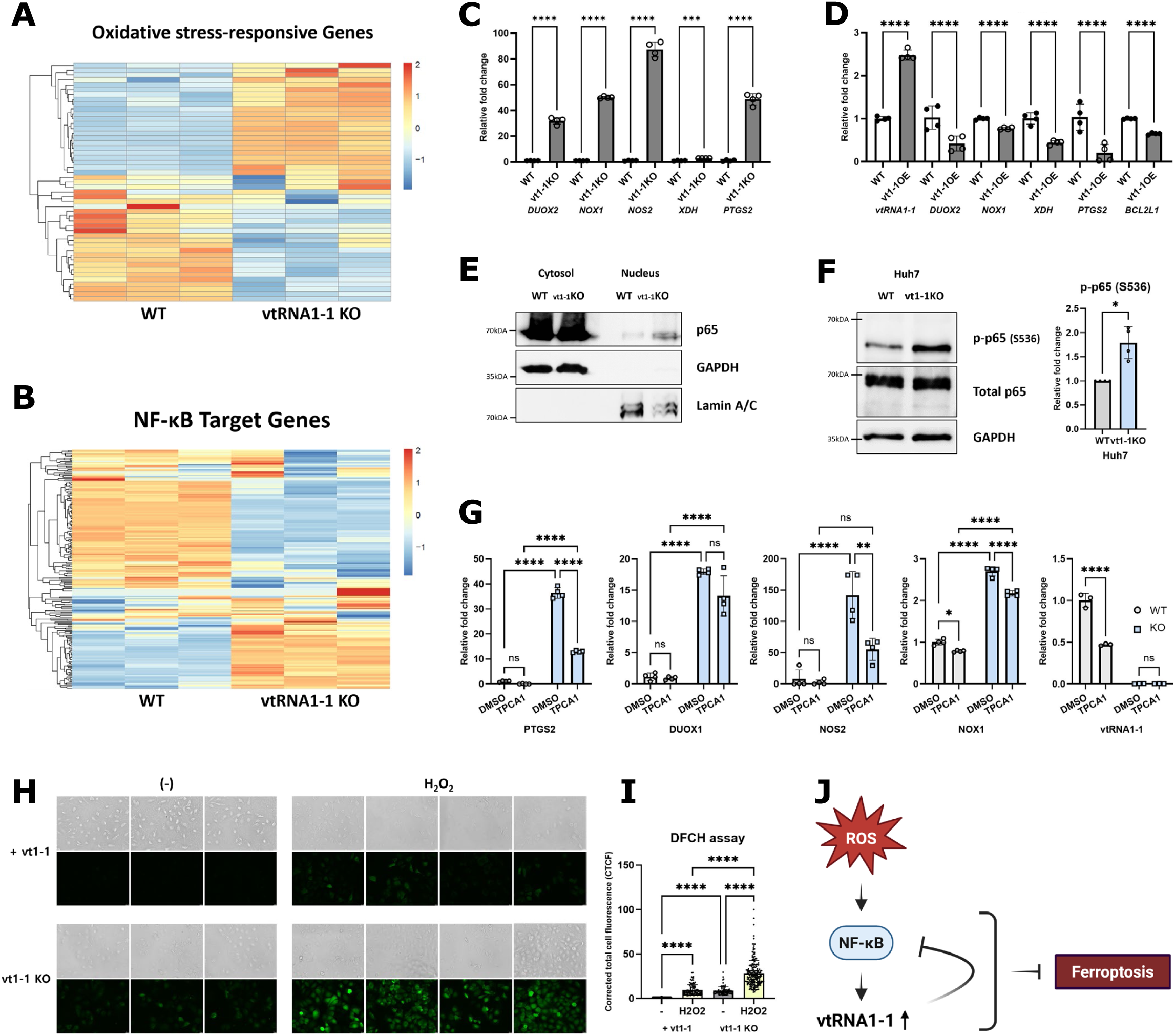
vtRNA1-1 suppresses NF-kB-mediated pro-oxidant expression and attenuates ROS accumulation. (A and B) Heatmaps showing the expression levels of oxidative stress-responsive genes (A) or NF-κB target genes (B) in Huh7 WT and vtRNA1-1 KO cells. (C and D) qPCR analyses of the indicated mRNAs and vtRNA1-1 in Huh7 WT and vtRNA1-1 KO (C) or vtRNA1-1 overexpression (D) cells (n=4). ****p<0.0001, ***p<0.001 (one-way ANOVA test) (E) Immunoblot analysis using the indicated antibodies. Following cytoplasm/nucleus fractionation, lysates were obtained from each fraction and analyzed. GAPDH and Lamin A/C served as cytoplasm and nucleus markers respectively. (F) Immunoblot analysis using the indicated antibodies. Lysates were obtained from each cell and analyzed. GAPDH served as a loading control. The intensity of phosphor-p65 was normalized to total p65 (right, n=4). Error bars indicate the standard deviation. *p<0.05 (t test) (G) qPCR analysis of the indicated mRNAs in Huh7 WT and vtRNA1-1 KO cells following treatment with TPCA-1 (10 μM) for 4 hours (n=4). ****p<0.0001, **p<0.01, ns: non-significant (two-way ANOVA test) (H) Intracellular ROS was measured by DCFH-DA assay. After treatment with 50 uM H_2_O_2_ for 30 minutes, vtRNA1-1 KO and vtRNA1-1-expressing (vtRNA1-1 +) Huh7 cells were incubated with 20 uM DCFH-DA for 30 minutes in the dark. Images were captured using a fluorescence microscope. (I) The fluorescent intensity shown in Fig. 6H was analyzed using Fiji software, and corrected total cell fluorescence (CTCF) was calculated. ****p<0.0001 (one-way ANOVA test) (J) A schematic illustration of proposed mechanisms by which vtRNA1-1 mediates ferroptosis resistance. Created in BioRender. Kong, EB. (2026)

To validate the functional role of this reciprocal relationship between NF-κB and vtRNA1-1 in regulating redox homeostasis, intracellular ROS levels were measured using a DCFH-DA probe. Intracellular ROS levels were elevated in vtRNA1-1 KO cells compared to vtRNA1-1 expressing cells **(Fig. 5H and 5I)**, indicating that the increased pro-oxidants by vtRNA1-1 deficiency induce ROS accumulation. To determine whether the loss of vtRNA1-1 also affects the ROS clearance system in cells, we further investigated the expression of antioxidant system-related genes. Although primary ROS scavenging enzymes, including SOD1, SOD2, GPX1, and CAT, showed no distinctive regulatory patterns in vtRNA1-1 KO cells **(Fig. S6C)**, the expression of antioxidants mediated by NRF2, a master transcriptional regulator of cellular redox homeostasis, was downregulated upon vtRNA1-1 depletion **(Fig. S6D and S6E)**. While the direct regulatory mechanism remains to be elucidated, our observations that vtRNA1-1 plays a key role in attenuating cellular ROS accumulation by modulating pivotal transcription factors, NF-κB and NRF2, align with their well-known antagonistic crosstalk [51,52]. Collectively, given that ROS accumulation, together with elevated ferrous ion concentration, is considered a major trigger of ferroptosis, the function of vtRNA1-1 in governing basal ROS level is critical to determining ferroptosis susceptibility in HCC cells.

### 3.6. vtRNA1-1-associated expression signature aligns with transcriptional programs of ferroptosis-related genes in the TCGA-LIHC cohort

We next assessed the clinical relevance of our findings by analyzing the TCGA-liver hepatocellular carcinoma (LIHC) cohort. To bypass the absence of vtRNA1-1 sequencing data in the TCGA database, we defined a vtRNA1-1-associated mRNA transcriptional signature by re-analyzing our previous transcriptomic data [36]. Compared with WT cells, the top 1,000 differentially expressed genes (DEGs) in vtRNA1-1 KO cells were defined as the vtRNA1-1-associated transcriptional signature. To investigate the relevance of vtRNA1-1 to hepatocarcinogenesis using this signature, principal component analysis (PCA) was performed across the entire TCGA-LIHC cohort, including both normal and tumor samples. The PCA demonstrated that tumor samples are clearly distinguishable from normal samples based on the vtRNA1-1 signature **(Fig. 6A)**. Additionally, hierarchical clustering analysis using this signature showed a distinct segregation between tumor and normal samples **(Fig. 6B)**, supporting our previous observations of elevated vtRNA1-1 in patient-derived tumor tissues [31].

**Figure 6.**
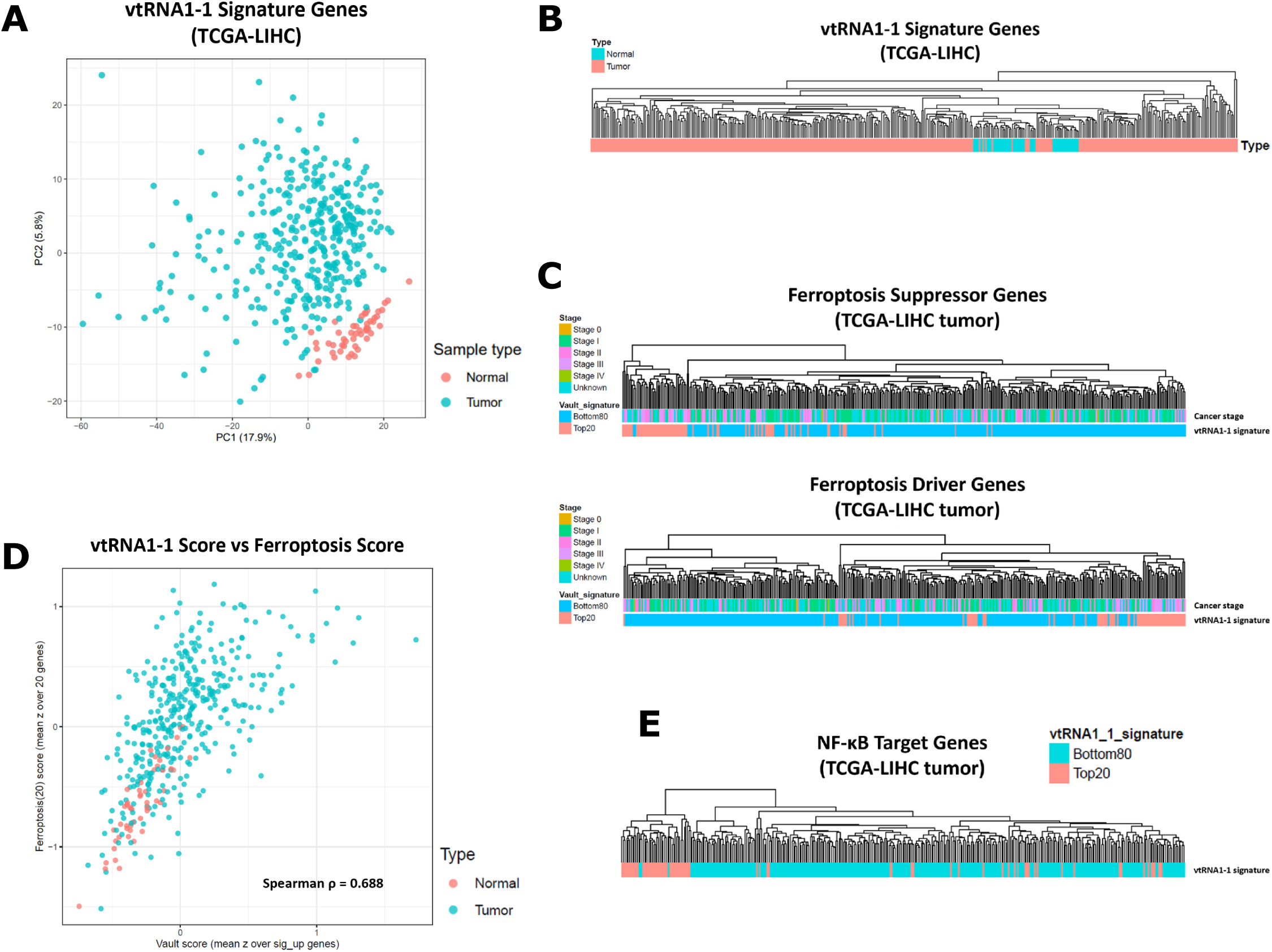
Clinical association of vtRNA1-1 expression with ferroptosis-related genes in HCC. (A) Principal component analysis (PCA) of the vtRNA1-1-associated signature genes showing entire normal (green) and tumor (red) samples in the TCGA-LIHC cohort. (B) Hierarchical clustering analysis using the vtRNA1-1-associated signature genes showing distinct clusters of normal (green) and tumor (red) samples in the TCGA-LIHC cohort. (C) Hierarchical clustering analyses using the ferroptosis-related genes (Suppressor: top, Driver: bottom) showing distinct clusters of Top20 (dark red) and Bottom80 (light pink) groups categorized by vtRNA1-1-associated signature score. (D) Correlation analysis between the vtRNA1-1-associated signature score and the expression of key ferroptosis genes showing a strong positive correlation (normal: green, tumor: red) (Spearman’s ρ=0.688) (E) Hierarchical clustering analysis using the NF-κB target genes showing distinct clusters of Top20 (dark red) and Bottom80 (light pink) groups categorized by vtRNA1-1-associated signature score.

Subsequently, to evaluate the clinical significance of vtRNA1-1 in the context of ferroptosis susceptibility in HCC cells, we divided the LIHC tumor samples into two groups based on the vtRNA1-1 signature score: the vtRNA1-1-high group (Top20) that showed relatively high signature scores, and the vtRNA1-1-low group (Bottom80) that expressed the vtRNA1-1 signature genes like vtRNA1-1 KO cells **(Fig. S7A)**. Clustering analysis and PCA based on ferroptosis-related genes sourced from the FerrDB [43], revealed distinct patterns between the Top20 and Bottom80 groups, indicating that the vtRNA1-1 signature correlated with transcriptional profile of ferroptosis-related genes without showing clear relevance to tumor stages **(Fig. 6C and S7B)**. A strong positive correlation (Spearman’s ρ =0.688) between the vtRNA1-1-associated signature score and the expression score of key ferroptosis genes corroborated these findings **(Fig. 6D)**. Additionally, the marked difference in the expression of NF-κB target genes, NRF2 target genes, and oxidative stress-responsive genes between Top20 and Bottom80 groups provided further evidence of vtRNA1-1 function in governing redox balance in a clinical context **(Fig. 6E, S7C and S7D)**. Notably, a marginal trend in overall survival difference between the two groups further suggests the potential relevance of vtRNA1-1 in HCC **(Fig. S7E)**. Altogether, given the apparent contributions of ferroptosis-related genes to clinical outcomes in HCC [53,54], these observations underscore a potential clinical importance of vtRNA1-1 in mediating ferroptosis susceptibility among HCC patients.

## 4. Discussion

In this study, we identified a previously unrecognized role of vtRNA1-1 in governing redox balance and ferroptosis susceptibility in human hepatocytes. Although the pro-survival non-coding vtRNA1-1 is known to regulate apoptosis in several types of cancer cell lines excluding HCC cells [31], its functional relevance to ferroptosis, a distinct type of ferrous ion-dependent regulated cell death (RCD), has remained entirely unexplored. Our findings demonstrate that vtRNA1-1 is induced by oxidative stress in an NF-κB dependent manner (**Fig.4**), and this short ncRNA suppresses NF-κB activity to decrease pro-oxidant expression, thereby hindering reactive oxygen species (ROS) accumulation (**Fig.5**). This reciprocal loop between vtRNA1-1 expression and NF-κB activation represents a pivotal mechanism for maintaining redox homeostasis, enhancing redox buffering capacity and ultimately determining the ferroptosis susceptibility in hepatocytes. Indeed, mining in the human TCGA-LIHC cohort database revealed markedly distinguishable vtRNA1-1-dependent transcriptional landscapes between liver tumor tissue and normal control tissue samples (**Fig. 6**).

In the context of severe stress, RCD constitutes a crucial process to preserve organismal integrity. Therefore, the determination of this irreversible option must be tightly, yet accurately regulated in a dynamic manner. Due to their dynamic nature, short ncRNAs are ideal for fine-tuning the RCD process in terms of kinetic plasticity with relatively low biosynthetic costs. Recently, roles of various short ncRNAs in regulating cell-fate determination have been uncovered [55]. As a pro-survival short ncRNA, vtRNA1-1 is elevated in response to stresses including viral infection and starvation, conferring survival benefits to cells and inhibiting apoptosis in cell lines originating from various cancer tissues [32,33,56,57] Here we identified that oxidative stress-induced vtRNA1-1 promotes hepatocyte viability by suppressing ferroptotic cell death. The oxidative stress, caused by imbalance between ROS production and antioxidant defense, results in significant dysfunctions in cells, including DNA damage, protein oxidation and lipid peroxidation. Given the role of the liver as a hub for metabolism and detoxification that generates a substantial amount of ROS, oxidative stress adaptation is particularly important for hepatocytes. Disrupted redox homeostasis in liver triggers oxidative stress, which in turn drives the pathogenesis of metabolic liver diseases such as alcoholic fatty liver disease, non-alcoholic fatty liver disease (NAFLD) and non-alcoholic steatohepatitis (NASH) [58,59]. Furthermore, chronic and severe oxidative stress accelerates the progression to fibrosis, cirrhosis, and eventually hepatocellular carcinoma (HCC) [60]. Considering the relevance of vtRNA1-1 in tumorigenesis [31,34], our new findings of oxidative stress-induced vtRNA1-1 expression may provide an unrecognized layer of understanding oxidative stress-driven HCC tumorigenesis. As oxidative stress is a critical driver of HCC, elevated vtRNA1-1 levels under oxidative stress suppresses cell death and potentially facilitates tumor progression. Thus, elucidating the connection between oxidative stress and the elevated expression of vtRNA1-1 in the context of HCC tumorigenesis is warranted.

Cells have an elaborated antioxidant system to maintain redox homeostasis. NRF2 is one of the master regulators of this system that controls the expression of protective genes against oxidative damage through binding to antioxidant responsive elements (ARE) [61,62]. We revealed that vtRNA1-1 is involved in the expression of NRF2 target genes such as HMOX1, GCLS, GSS, GSR, and TXRND1, whose expressions were downregulated in vtRNA1-1 KO cells (**Fig.S6**). Another key player in governing redox balance is the transcription factor NF-κB. Activated by excessive ROS, NF-κB regulates both pro- and antioxidant genes expression in a context-dependent manner. We demonstrated the reciprocal regulation between vtRNA1-1 and NF-κB to maintain redox homeostasis. vtRNA1-1 depletion induces pro-oxidants, including DUOX2, NOS2, NOX1, XDH, and PTGS2 to increase ROS accumulation, whereas the reintroduction of vtRNA1-1 or NF-κB inhibition rescued the elevated expression of these genes in KO cells (**Figs.5**). Collectively, vtRNA1-1 regulates redox balance by modulating two key transcription factors, NRF2 and NF-κB. Considering the importance of these transcription factors and the complexity of their relationship [61,63,64], this study provides new insights not only into the understanding of HCC progression, but also unveils a so far hidden crosstalk between NRF2 and NF-κB.

Despite these unprecedented observations, it is important to note that the precise underlying molecular details through which vtRNA1-1 regulates two key transcription factors remain to be elucidated. Given that the vast majority of vtRNAs do not associate with vault particles [57,65], discovering novel interacting proteins of vtRNA1-1 can be a promising approach to uncover the underlying mechanisms. Indeed, studies have demonstrated that vtRNAs regulate cellular processes by interacting with cytoplasmic proteins. For instance, vtRNA1-1 interacts with the CUG-binding protein 1 (CUGBP1) to inhibit the expression of claudin-1 and occludin in intestinal epithelial cells [66]. In Huh7 cells, vtRNA1-1 directly binds to p62 using their central region to regulate p62-dependent autophagy [56]. In addition, TRIM21 and TRIM25 have been recently demonstrated in HCC cells to modulate vtRNA1-1 stability [34]. Additionally, mouse vtRNA (mvtRNA) exhibits direct binding to MEK1 and stimulates MAPK signaling for synaptogenesis [67]. Recently, Stok et al. verified that vtRNA1 paralogs interact with ELAVL1 and hnRNP C and facilitate their export from the nucleus [68]. Given the exclusive function of vtRNA1-1 in regulating redox homeostasis among the three vtRNA1 paralogs (**Fig.4 and 5**), hnRNP U, a distinct binding partner of vtRNA1-1 in the proteomic screening by Stok et al. is worthy for further investigation. The very recently proposed role of hnRNP U in regulating ferroptosis in colon adenocarcinoma makes this putative protein/vtRNA1-1 interaction an attractive target for further studies [69].

Through comparative analysis, we observed that the expression pattern of vtRNA1 paralogs varies markedly across the different liver cell lines **(Fig. 1D)**. Among them, we focused on vtRNA1-1 that shows a strong negative correlation with ferroptosis sensitivity. Cells expressing high levels of vtRNA1-1 display a ferroptosis-resistant phenotype, whereas ferroptosis-sensitive cells express relatively low vtRNA1-1 **(Fig. 1F)**. These findings suggest vtRNA1-1 as a potential biomarker for predicting ferroptosis susceptibility in liver cancer. Accordingly, the assessment of vtRNA1-1 in liver cancer tissue could be used to determine the suitability of ferroptosis-based therapeutic strategies. Furthermore, we revealed that vtRNA1-1 depletion sensitizes HCC cells to ferroptosis, while overexpression of vtRNA1-1 confers ferroptosis resistance in liver cells **(Fig.2)**. These results therefore suggest additional avenues in which this short ncRNA can be utilized not only as a biomarker, but also as an effective target for anti-liver cancer strategies. Given the elevated vtRNA1-1 expression in patient-derived liver cancer tissues [31], targeting vtRNA1-1 to enhance the efficacy of ferroptosis-based HCC therapy appears attractive.

To summarize, this study identified a novel role of vtRNA1-1 in HCC cells by governing the redox balance, thereby determining ferroptosis susceptibility. Elevated vtRNA1-1 expression observed in highly ferroptosis resistant HCC cell lines negatively correlates with ferroptosis sensitivity. We further uncovered oxidative stress-induced vtRNA1-1 expression in an NF-κB dependent manners, consequently establishing a reciprocal loop with NF-κB to suppress pro-oxidant gene expression and ROS accumulation. As a pro-survival ncRNA, vtRNA1-1 attenuates cellular ROS accumulation, conferring HCC cells resistance to oxidative stress and ferroptosis. Our investigation establishes vtRNA1-1 as a potential therapeutic target for overcoming ferroptosis resistance in liver cancer.

## Acknowledgement

We would like to thank Maximilian Wolfensberger and Valentina Pecoraro for technical assistance and fruitful discussions, respectively. Our thanks are extended to Adrian Keogh for providing hepatocyte cell lines (IHH, HepG2, HepG2-C3A, Hep3B, SNU423, SNU449, and SNU475). The graphical abstract, Figure 3A and Figure 5J were made using BioRender.

## Author contributions

EB.K. performed the majority of the experiments and wrote the first draft of the manuscript. C.O.R. and A.S. contributed to Figures 3 and Figure 5 respectively. D.S.T. performed RNAseq and TCGA cohort analyses (under the supervision of D.S.). N.P. conceived the project and supervised the study (together with D.S.). All authors contributed to data interpretation, analysis and commented on the manuscript. Funding was acquired by N.P.

