## Supplemental Tables and Figures for "Stress-induced vtRNA1-1 modulates redox homeostasis and ferroptosis susceptibility in hepatocellular carcinoma cells"

**Probe sequences for northern blot**

| Probe (NB) | Sequence (5' - 3') |
|---|---|
| vtRNA1-1 | GCTTGTTTCAATTAAAGAACTGTCG |
| vtRNA1-2 | AGGTGGTTACAATGTACTCGAAG |
| vtRNA1-3 | GAGGTGGTTTGATGACACGCGAA |
| 5.8S rRNA | TCCTGCAATTCACATTAATTCTCGAGCTAGC |

**Primer sequences for RT-qPCR**

| Primer (qPCR) | Sequence (5' - 3') |
|---|---|
| vtRNA1-1 (Fw) | GGCTGGCTTTAGCTCAGC |
| vtRNA1-1 (Rv) | CCAGACAGGTTGCTTGTT |
| GAPDH (Fw) | TCAAGGCTGAGAACGGGAAG |
| GAPDH (Rv) | CGCCCCACTTGATTTTGGAG |
| 18S rRNA (Fw) | GAGAAACGGCTACCACATCCA |
| 18S rRNA (RV) | CTCCAATGGATCCTCGTTAAAGG |
| CHAC1 (Fw) | GGTGACGCTCCTTGAAGATC |
| CHAC1 (Rv) | CACTGCCTCTCGCACATT |
| ALOX15 (Fw) | CTTCCTAACCTACAGCTCCTTCT |
| ALOX15 (Rv) | ACAGCCACGTCTGTCTTATAGT |
| PTGS2 (Fw) | GATGATTGCCCGACTCCCTT |
| PTGS2 (Rv) | GGCCCTCGCTTATGATCTGT |
| DUOX2 (Fw) | ACGCAGCTCTGTGTCAAAGGT |
| DUOX2 (Rv) | TGATGAACGAGACTCGACAGC |
| NOX1 (Fw) | CAATCTCTCTCCTGGAATGGCATC |
| NOX1 (Rv) | CTGCTGCTCGGATATGAATGGAG |
| NOS2 (Fw) | TGCAGACACGTGCGTTACTC |
| NOS2 (Rv) | GGTAGCCAGCATAGCGGATG |
| XDH (Fw) | GACAGTTGTGGCTCTTGAGGT |
| XDH (Rv) | GGAAGGTTGGTTTTGCACAGC |
| BCL2L1 (Fw) | CGGTACCGGCGGGCATTCAG |
| BCL2L1 (Rv) | CGGCTCTCGGCTGCTGCATT |
| FSP1 (Fw) | CCCTGCCCTTCTCTCATCTTA |
| FSP1 (Rv) | TGTCCTCATAGGCCTGGATAG |
| SOD1 (Fw) | CCAGTGCAGGTCCTCACTTTA |
| SOD1 (Rv) | GTCTCCAACATGCCTCTCTTCA |
| SOD2 (Fw) | GGCTCCGGTTTTGGGGTATC |
| SOD2 (Rv) | GCATGATCTGCGCGTTGATG |
| GPX1 (Fw) | GTGTATGCCTTCTCGGCGC |
| GPX1 (Rv) | GACCGTGGTGCCTCAGAG |
| CAT (Fw) | CAGCGACCAGATGCAGCA |
| CAT (Rv) | CCACGGGGCCCTACTGTAA |
| HMOX1 (Fw) | AAGACTGCGTTCCTGCTCAA |
| HMOX1 (Rv) | GGGCAGAATCTTGCACTTTGT |
| GSR (Fw) | ATGGTCTGTGCTAACAAGGAAGA |

| | |
|---|---|
| GSR (Rv) | GTTGCTCCCATCTTCACTGCA |
| GSS (Fw) | GGGAGCCTCTTGCAGGATAAA |
| GSS (Rv) | GAATGGGGCATAGCTCACCAC |
| GCLC (Fw) | GGAGACCAGAGTATGGGAGTT |
| GCLC (Rv) | CCGGCGTTTTCGCATGTTG |
| TXRND1 (Fw) | TCACCCCAGTTGCAATCC |
| TXRND1 (Rv) | GTTGGAACATTTTCATAGTCACA |
| TXRND2 (Fw) | CGGCTTCGACCAGCAAATG |
| TXRND2 (Rv) | ACAGGACGGTGTCAAAGGTG |

### Figure S1

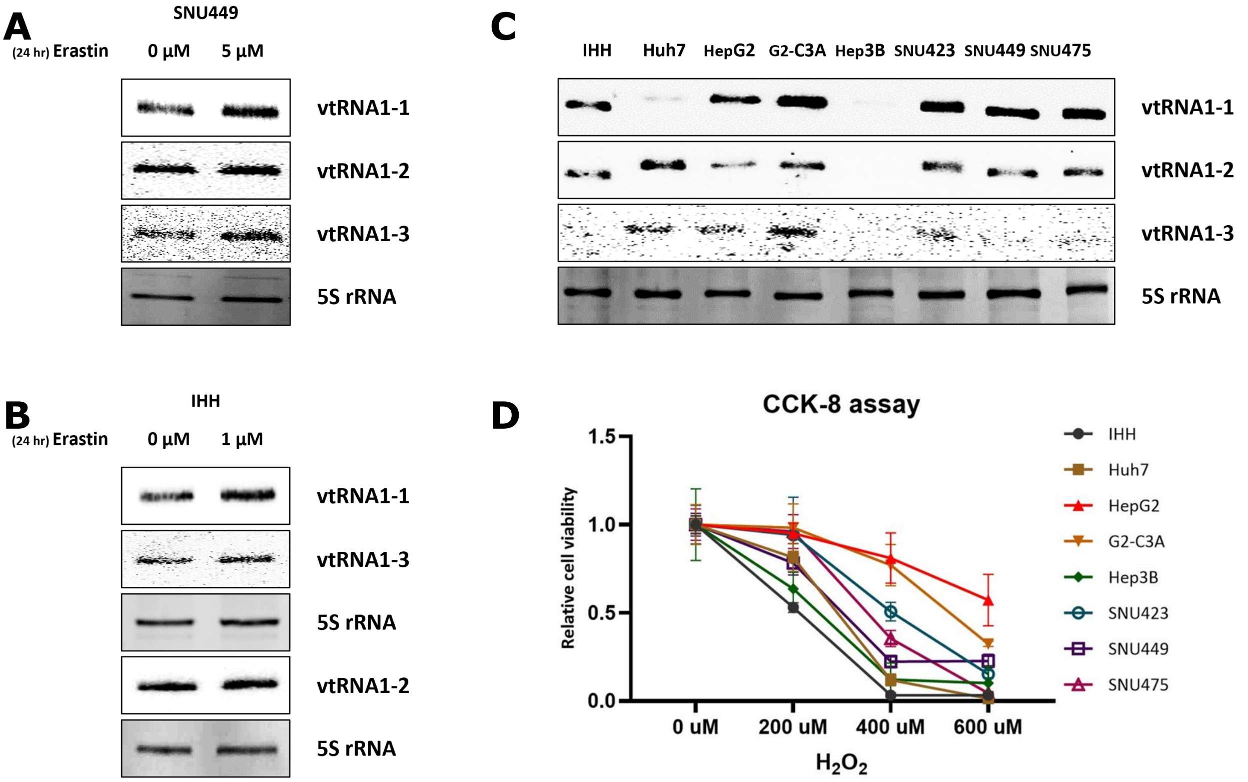

A
SNU449
(24 hr) Erastin
0 µM
5 µM
vtRNA1-1
vtRNA1-2
vtRNA1-3
5S rRNA
B
IHH
(24 hr) Erastin
0 µM
1 µM
vtRNA1-1
vtRNA1-3
5S rRNA
vtRNA1-2
5S rRNA
C
IHH
Huh7
HepG2
G2-C3A
Hep3B
SNU423
SNU449
SNU475
vtRNA1-1
vtRNA1-2
vtRNA1-3
5S rRNA
D
CCK-8 assay
Relative cell viability
1.5
1.0
0.5
0.0
0 uM
200 uM
400 uM
600 uM
H2O2
IHH
Huh7
HepG2
G2-C3A
Hep3B
SNU423
SNU449
SNU475

### Figure S2

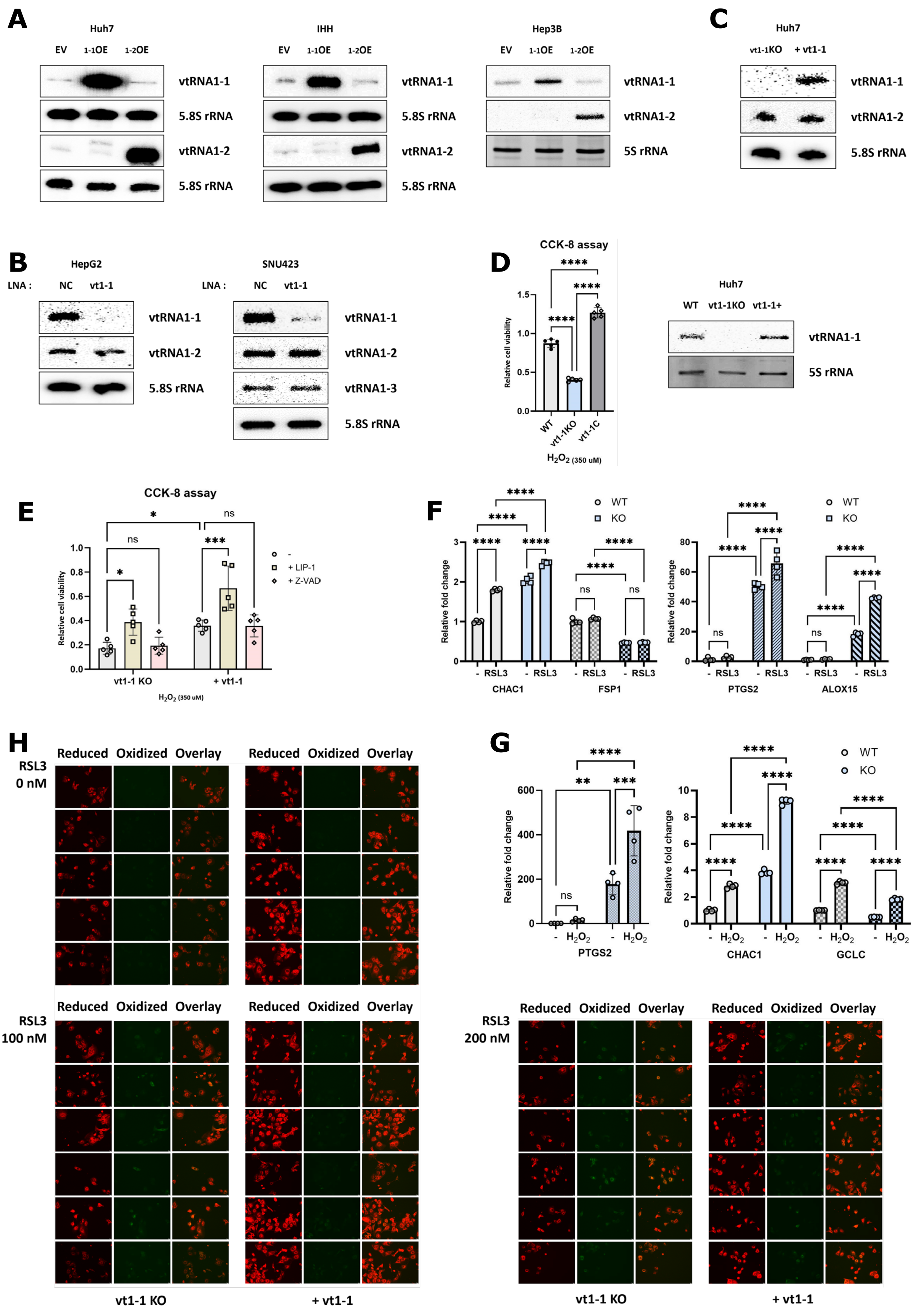
A
Huh7
EV 1-1OE 1-2OE
vtRNA1-1
5.8S rRNA
vtRNA1-2
5.8S rRNA
IHH
Hep3B
vtRNA1-2
5S rRNA
C
Huh7
vt1-1KO + vt1-1
vtRNA1-1
vtRNA1-2
5.8S rRNA
B
HepG2
SNU423
LNA : NC vt1-1
vtRNA1-3
D
CCK-8 assay
Relative cell viability
WT vt1-1KO vt1-1C
H2O2 (350 uM)
Huh7
WT vt1-1KO vt1-1+
E
CCK-8 assay
- + LIP-1 + Z-VAD
vt1-1 KO + vt1-1
F
WT KO
Relative fold change
- RSL3
CHAC1 FSP1 PTGS2 ALOX15
G
- H2O2
PTGS2 CHAC1 GCLC
H
Reduced Oxidized Overlay
RSL3 0 nM
RSL3 100 nM
RSL3 200 nM
vt1-1 KO
+ vt1-1

### Figure S3

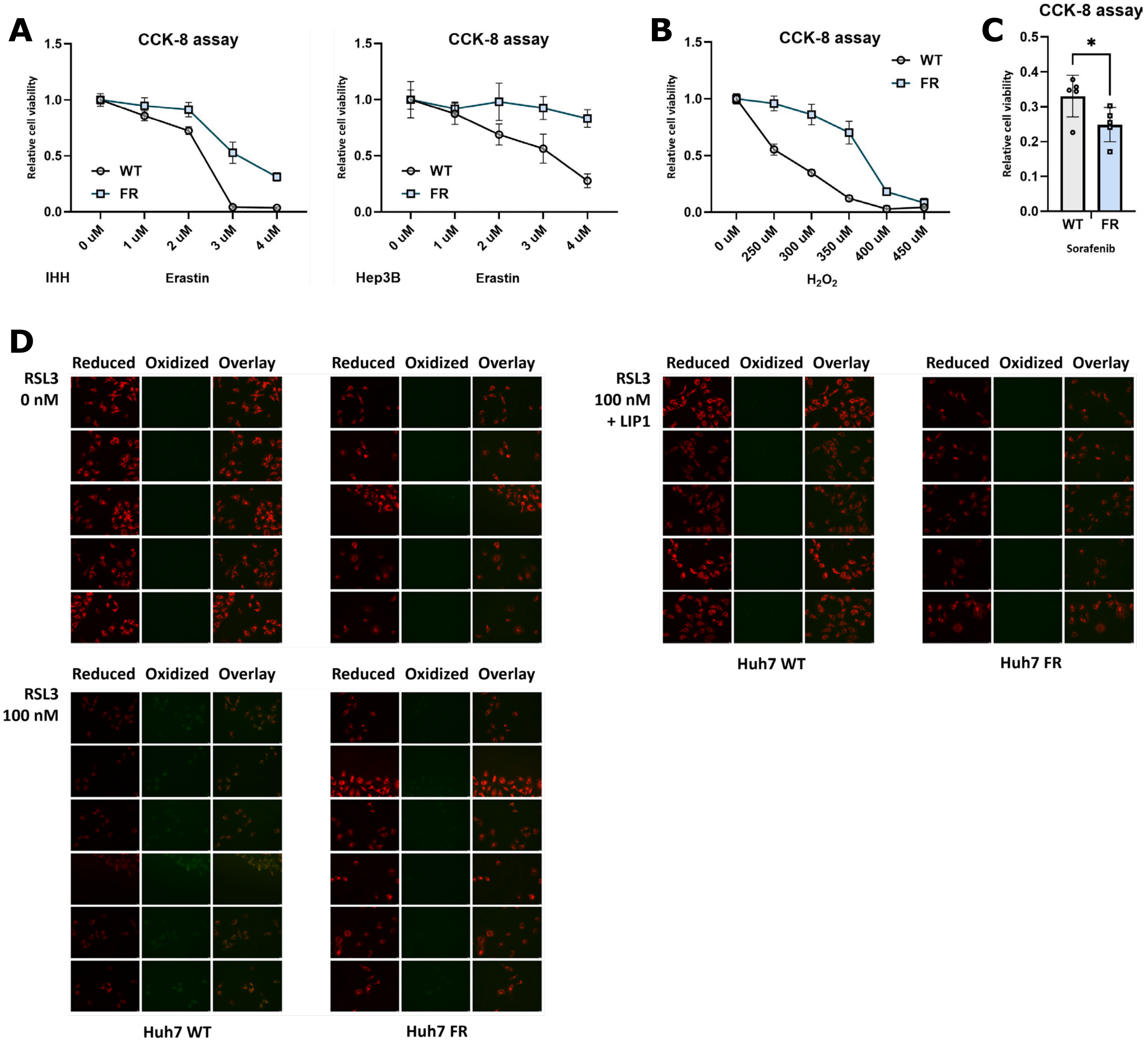

A
CCK-8 assay
Relative cell viability
1.5
1.0
0.5
0.0
WT
FR
0 uM
1 uM
2 uM
3 uM
4 uM
IHH
Erastin
CCK-8 assay
Hep3B
Erastin
B
CCK-8 assay
0 uM
250 uM
300 uM
350 uM
400 uM
450 uM
$H_2O_2$
C
CCK-8 assay
*
0.5
0.4
0.3
0.2
0.1
0.0
Sorafenib
D
Reduced
Oxidized
Overlay
RSL3
0 nM
RSL3
100 nM
+ LIP1
RSL3
100 nM
Huh7 WT
Huh7 FR

### Figure S4

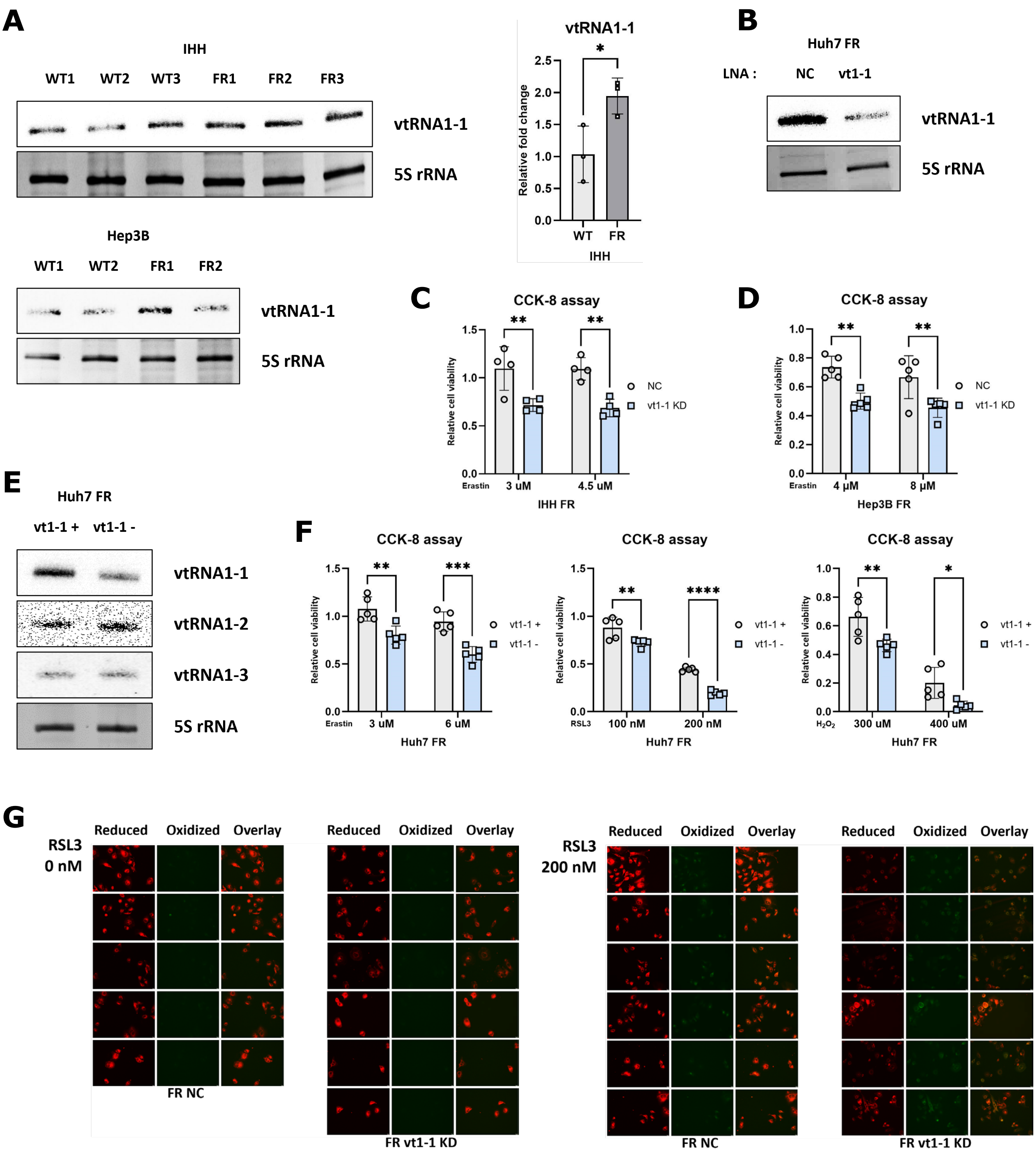

A
IHH
WT1 WT2 WT3 FR1 FR2 FR3
vtRNA1-1
5S rRNA
vtRNA1-1
Relative fold change
WT FR
IHH
Hep3B
WT1 WT2 FR1 FR2
vtRNA1-1
5S rRNA
B
Huh7 FR
LNA : NC vt1-1
vtRNA1-1
5S rRNA
C
CCK-8 assay
Relative cell viability
NC
vt1-1 KD
Erastin 3 uM 4.5 uM
IHH FR
D
CCK-8 assay
Relative cell viability
NC
vt1-1 KD
Erastin 4 μM 8 μM
Hep3B FR
E
Huh7 FR
vt1-1 + vt1-1 -
vtRNA1-1
vtRNA1-2
vtRNA1-3
5S rRNA
F
CCK-8 assay
Relative cell viability
vt1-1 +
vt1-1 -
Erastin 3 uM 6 uM
Huh7 FR
CCK-8 assay
RSL3 100 nM 200 nM
Huh7 FR
CCK-8 assay
H₂O₂ 300 uM 400 uM
Huh7 FR
G
RSL3 0 nM
Reduced Oxidized Overlay
FR NC
FR vt1-1 KD
RSL3 200 nM
FR NC
FR vt1-1 KD

### Figure S5

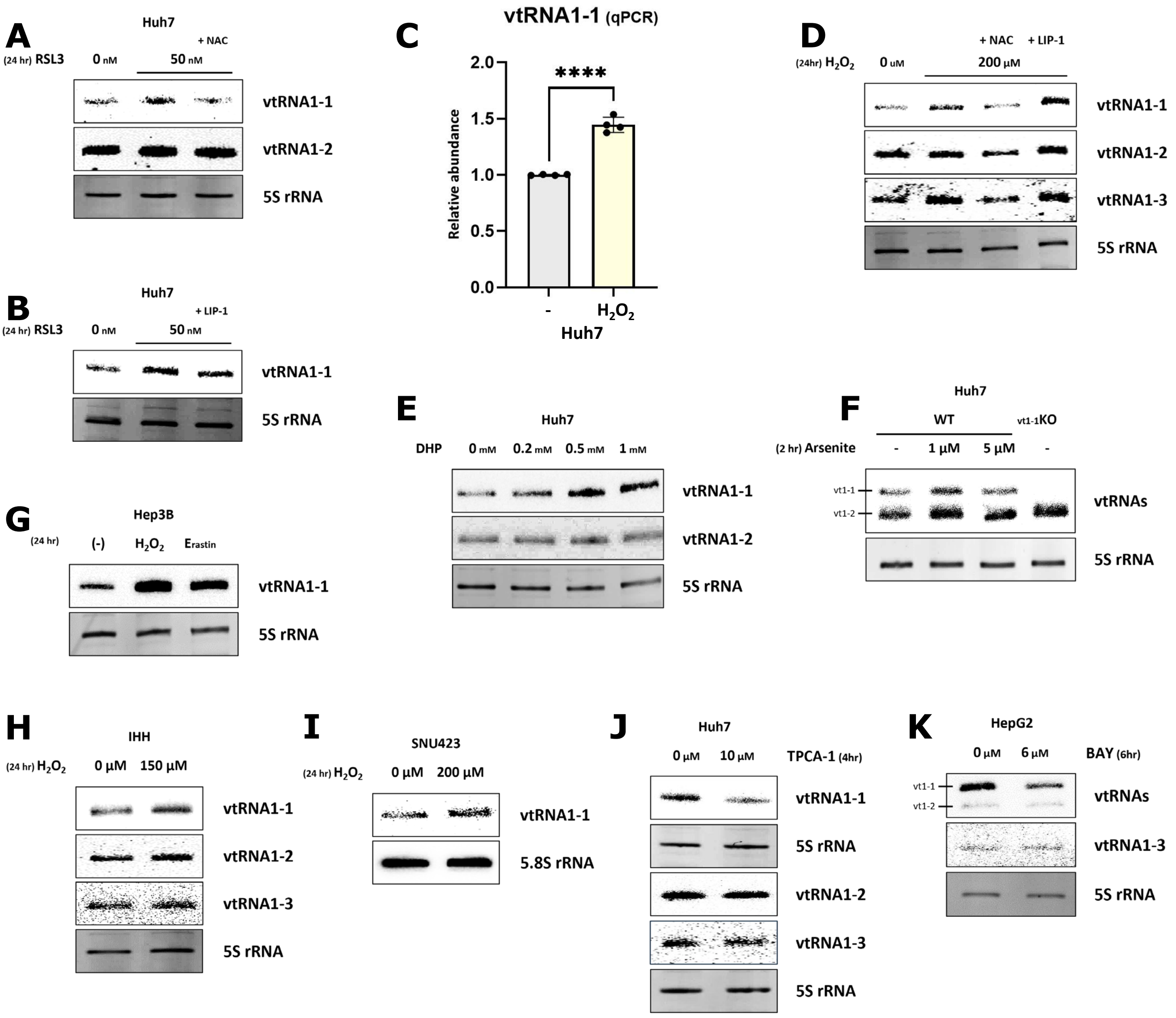

A
Huh7
(24 hr) RSL3
0 nM
50 nM
+ NAC
vtRNA1-1
vtRNA1-2
5S rRNA
B
Huh7
(24 hr) RSL3
0 nM
50 nM
+ LIP-1
vtRNA1-1
5S rRNA
C
vtRNA1-1 (qPCR)
****
Relative abundance
2.0
1.5
1.0
0.5
0.0
-
$H_2O_2$
Huh7
D
(24hr) $H_2O_2$
0 uM
200 μM
+ NAC
+ LIP-1
vtRNA1-1
vtRNA1-2
vtRNA1-3
5S rRNA
E
Huh7
DHP
0 mM
0.2 mM
0.5 mM
1 mM
vtRNA1-1
vtRNA1-2
5S rRNA
F
Huh7
WT
vt1-1KO
(2 hr) Arsenite
-
1 μM
5 μM
-
vt1-1
vt1-2
vtRNAs
5S rRNA
G
Hep3B
(24 hr)
(-)
$H_2O_2$
Erastin
vtRNA1-1
5S rRNA
H
IHH
(24 hr) $H_2O_2$
0 μM
150 μM
vtRNA1-1
vtRNA1-2
vtRNA1-3
5S rRNA
I
SNU423
(24 hr) $H_2O_2$
0 μM
200 μM
vtRNA1-1
5.8S rRNA
J
Huh7
0 μM
10 μM
TPCA-1 (4hr)
vtRNA1-1
5S rRNA
vtRNA1-2
vtRNA1-3
5S rRNA
K
HepG2
0 μM
6 μM
BAY (6hr)
vt1-1
vt1-2
vtRNAs
vtRNA1-3
5S rRNA

### Figure S6

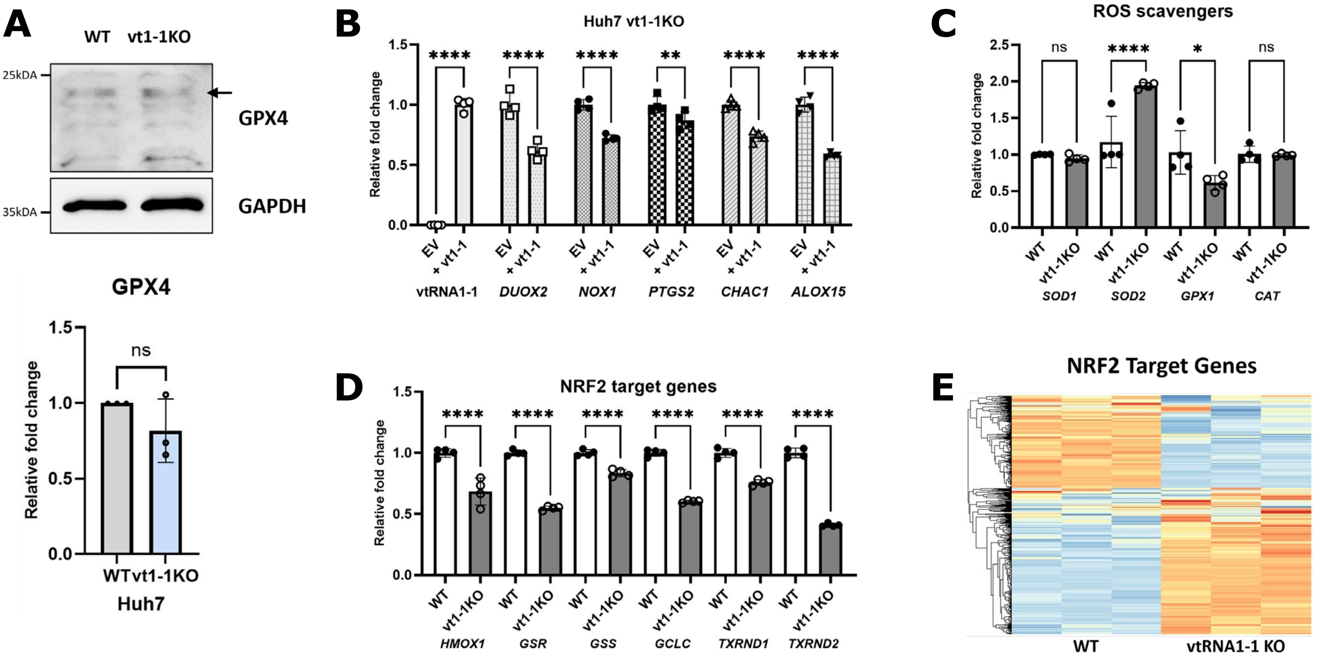

A
WT vt1-1KO
25kDA
35kDA
GPX4
GAPDH
GPX4
Relative fold change
ns
WT vt1-1KO
Huh7
B
Huh7 vt1-1KO
Relative fold change
EV
+ vt1-1
vtRNA1-1
DUOX2
NOX1
PTGS2
CHAC1
ALOX15
C
ROS scavengers
Relative fold change
WT
vt1-1KO
SOD1
SOD2
GPX1
CAT
D
NRF2 target genes
Relative fold change
HMOX1
GSR
GSS
GCLC
TXRND1
TXRND2
E
NRF2 Target Genes
WT
vtRNA1-1 KO

### Figure S7

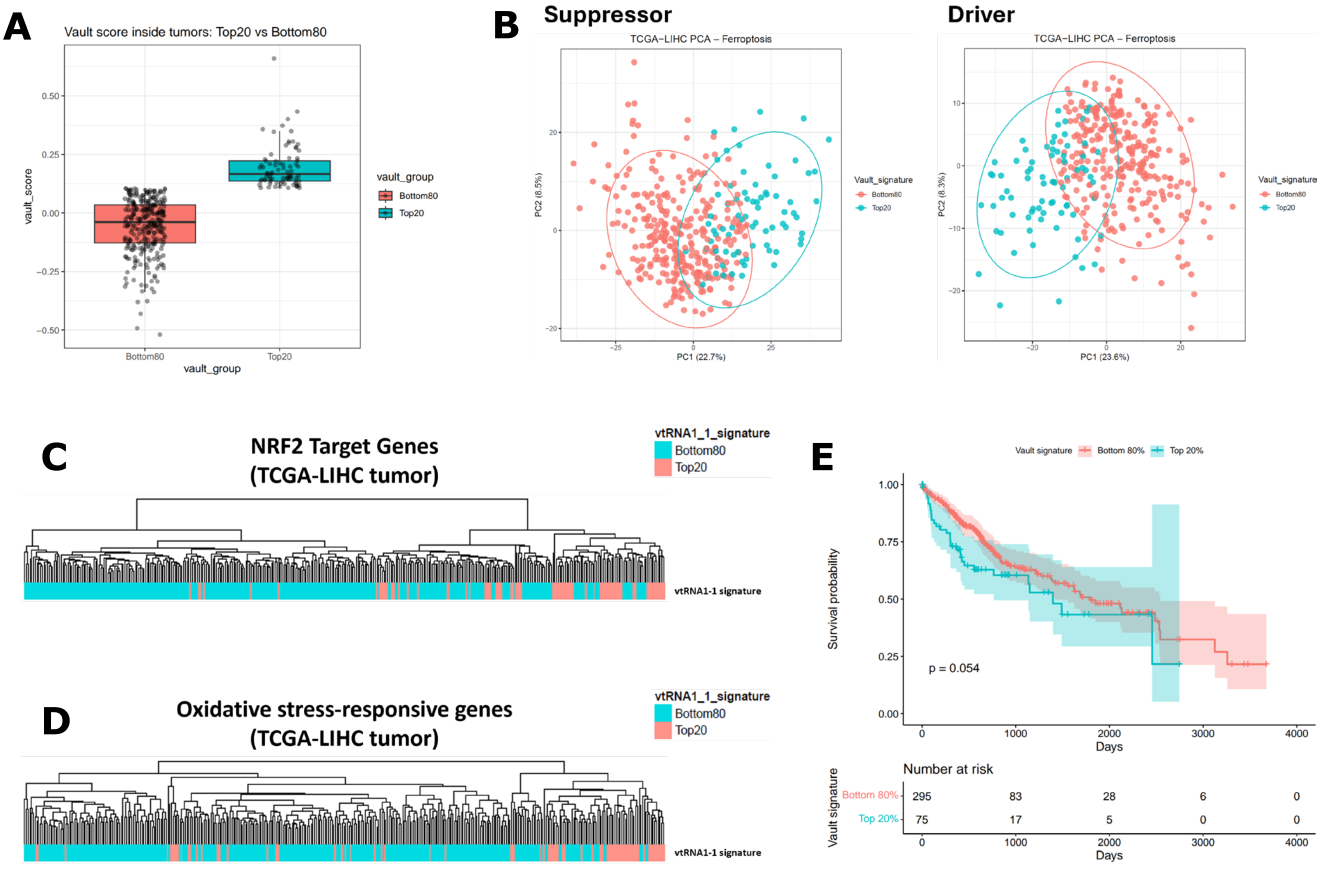

A
Vault score inside tumors: Top20 vs Bottom80
vault_score
vault_group
Bottom80
Top20
B
Suppressor
Driver
TCGA−LIHC PCA – Ferroptosis
PC1 (22.7%)
PC2 (8.5%)
PC1 (23.6%)
PC2 (8.3%)
Vault_signature
C
NRF2 Target Genes
(TCGA-LIHC tumor)
vtRNA1_1_signature
vtRNA1-1 signature
D
Oxidative stress-responsive genes
(TCGA-LIHC tumor)
E
Vault signature
Bottom 80%
Top 20%
Survival probability
p = 0.054
Days
Number at risk
295
83
28
6
0
75
17
5
0
0

**Figure S1.** (A and B) Northern blot analyses of vtRNAs. Total RNA was extracted from indicated cells treated with erastin for 24 hours and analyzed. 5S rRNA served as a loading control. (C) Northern blot analysis of vtRNAs. Total RNA was extracted from eight human liver cell lines and analyzed. 5S rRNA served as a loading control. (D) The indicated cells were treated with $H_2O_2$ for 24 hours and the cell viability was measured by CCK-8 assay.

**Figure S2.** (A and B) Northern blot analyses of vtRNAs. Following vtRNA1-1 or vtRNA1-2 overexpression (A) or LNA-mediated vtRNA1-1 knockdown (B), total RNA was extracted and analyzed. 5.8S rRNA and 5S rRNA served as loading controls. EV denotes empty vector and NC denotes negative control (mock transfection). (C) Northern blot analysis of vtRNAs. Following reintroduction of vtRNA1-1 into vtRNA1-1 KO Huh7 cells, total RNA was extracted and analyzed. 5.8S rRNA served as a loading control. (D) The indicated cells were treated with $H_2O_2$ for 24 hours, and the relative cell viability was measured by CCK-8 assay (left), normalized to the untreated cells. Northern blot analysis of vtRNA1-1. Total RNA was extracted from the indicated cells and analyzed. 5S rRNA served as a loading control. (right) (E) The indicated cells were treated with $H_2O_2$ and indicated inhibitors for 24 hours, and the relative cell viability was measured by CCK-8 assay, normalized to the untreated cells. (F and G) qPCR analyses of the indicated mRNAs in Huh7 WT and vtRNA1-1 KO cells treated with RSL3 (F) or $H_2O_2$ (G) (n=4). (H) Lipid peroxidation was measured using C11-BODIPY fluorescent probe. After pre-treatment with RSL3, vtRNA1-1 KO and vtRNA1-1-expressing (vtRNA1-1 +) Huh7 cells were incubated with 10 uM of BODIPY 581/591 for 30 minutes in the dark. Images were captured using a fluorescence microscope. ****$p<0.0001$, *** $p<0.001$, **$p<0.01$, ns: non-significant (two-way ANOVA test)

**Figure S3.** (A-C) The indicated cells were treated with erastin (A), $H_2O_2$ (B) or sorafenib (C) for 24 hours and the cell viability was measured by CCK-8 assay. *$p<0.05$ (t-test) (D) Lipid peroxidation was measured using C11-BODIPY fluorescent probe. After pre-treatment with RSL3 and Lip-1, Huh7 WT and ferroptosis-resistant (FR) cells were incubated with 10 uM of BODIPY 581/591 for 30 minutes in the dark. Images were captured using a fluorescence microscope.

**Figure S4.** (A) Northern blot analysis of vtRNAs. Total RNA was extracted from WT and FR cells and analyzed. 5S rRNA served as a loading control. The intensity of vtRNA1-1 in IHH was normalized to 5S rRNA (left, n=3). Error bars indicate the standard deviation. *$p<0.05$ (t-test) (B) Northern blot analysis of vtRNAs. Following LNA-mediated vtRNA1-1 knockdown, total RNA was extracted and analyzed. 5S rRNA served as loading controls. NC denotes negative control (mock transfection). (C and D) Following LNA-mediated knockdown of vtRNA1-1, indicated FR cells were treated with erastin for 24 hours, and the cell viability was measured by CCK-8 assay, normalized to the untreated cells. (E) Northern blot analysis of vtRNAs. Total RNA was extracted from control and CRISPR/Cas9-mediated vtRNA1-1-depleted FR cells and analyzed. 5S rRNA served as a loading control. (F) Following CRISPR/Cas9-mediated depletion of vtRNA1-1, Huh7 FR cells were treated with ferroptosis inducer for 24 hours, and the cell viability was measured by CCK-8 assay, normalized to the untreated cells. (G) Lipid peroxidation was measured using C11-BODIPY fluorescent probe. After pre-treatment with RSL3, control and LNA-mediated vtRNA1-1-depleted Huh7 FR cells were incubated with 10 uM of BODIPY 581/591 for 30 minutes in the dark. Images were captured using a fluorescence microscope. ****$p<0.0001$, *** $p<0.001$, **$p<0.01$, *$p<0.05$, ns: non-significant (two-way ANOVA test)

**Figure S5.** (A-B and D-K) Northern blot analyses of vtRNAs. Total RNA was extracted from cells treated with indicated drugs and analyzed. 5.8S rRNA or 5S rRNA served as loading controls. Final concentrations were as follows: 5 mM NAC, 1 μM LIP-1, and 10 μM TPCA-1. (C) qPCR analysis of vtRNA1-1 in Huh7 WT cells treated $H_2O_2$ (G) (n=4). ****$p<0.0001$ (t-test)

**Figure S6.** (A) Immunoblot analysis using the indicated antibodies. Lysates were obtained from each cell and analyzed. GAPDH served as a loading control. The intensity of GPX4 was normalized to GAPDH (right, n=3). Error bars indicate the standard deviation. ns: non-significant (t test) (B-D) qPCR analyses of the indicated mRNAs in Control and vtRNA1-1-overexpressed Huh7 cells (B) or Huh7 WT and vtRNA1-1 KO cells (C and D) (n=4). EV denotes empty vector (mock transfection). (E) A heatmap showing the expression levels of NRF2 target genes in Huh7 WT and vtRNA1-1 KO cells. ****$p<0.0001$, **$p<0.01$, ns: non-significant (two-way ANOVA test)

**Figure S7.** (A) Box plot of Top20 (dark red) and Bottom80 (light pink) groups in the TCGA-LIHC tumor cohorts, stratified by vtRNA1-1-associated signature score. (B) Principal component analysis (PCA) of the ferroptosis-related genes (Suppressor: left, Driver: right) showing Top20 (dark red) and Bottom80 (light pink) groups in the TCGA-LIHC tumor cohorts. (C and D) Hierarchical clustering analysis using the NRF2 target genes (C) or oxidative stress-responsive genes (D) showing distinct clusters of Top20 (dark red) and Bottom80 (light pink) groups categorized by vtRNA1-1-associated signature score. (E) Survival analysis of TCGA-LIHC cohorts based on the vtRNA1-1-associated signature score. (Top20: dark red, Bottom80: light pink) (p value=0.054)
